# Foxj1 controls olfactory ciliogenesis and differentiation program of the olfactory sensory neurons

**DOI:** 10.1101/2023.05.10.540158

**Authors:** Dheeraj Rayamajhi, Mert Ege, Kirill Ukhanov, Christa Ringers, Yiliu Zhang, Inyoung Jeong, Percival P. D’Gama, Summer Shijia Li, Mehmet Ilyas Cosacak, Caghan Kizil, Hae-Chul Park, Emre Yaksi, Jeffrey R. Martens, Steven L. Brody, Nathalie Jurisch-Yaksi, Sudipto Roy

## Abstract

In vertebrates, olfactory receptors localize on multiple cilia elaborated on dendritic knobs of olfactory sensory neurons (OSNs). Although olfactory cilia dysfunction can cause anosmia, how their differentiation is programmed at the transcriptional level has remained largely unexplored. We discovered in zebrafish and mice that Foxj1, a forkhead domain-containing transcription factor linked with motile cilia biogenesis, is expressed in OSNs and required for olfactory epithelium (OE) formation. In keeping with the immotile nature of olfactory cilia, we observed that ciliary motility genes are repressed in zebrafish, mouse, and human OSNs. Strikingly, we also found that besides ciliogenesis, Foxj1 controls the differentiation of the OSNs themselves by regulating their cell type-specific gene expression, such as that of *olfactory marker protein* (*omp*) involved in odor-evoked signal transduction. In line with this, response to bile acid, an odor detected by OMP-positive OSNs, was significantly diminished in *foxj1* mutant zebrafish. Taken together, our findings establish how the canonical Foxj1-mediated motile ciliogenic transcriptional program has been repurposed for the biogenesis of immotile olfactory cilia and for development of the OSNs.

## INTRODUCTION

Olfaction plays important roles in food and mate choice, and also in the avoidance of predators, making it a vital sensory modality for preservation and reproduction. The olfactory system consists of a highly specialized epithelium within the nasal cavity that contains odor-responsive neurons called OSNs. The OSNs have apical specializations called dendritic knobs, which sprout multiple long cilia that intertwine to form an expanded receptive surface, embedded in nasal mucus (Menco and Farbman 1985, Ache and Young 2005). This ciliary meshwork is the site for the localization of olfactory receptors, which on binding to odors, activate signaling that is then transmitted to the olfactory bulb (OB) in the brain for perception (Ache and Young 2005, McClintock, Khan et al. 2020). In humans, the loss of smell is a characteristic feature of many ciliary disorders (Kulaga, Leitch et al. 2004, Iannaccone, Mykytyn et al. 2005, Jenkins, McEwen et al. 2009). Consistent with this, mutation of several ciliary genes in model organisms, like the zebrafish and mouse, have been shown to impair olfactory cilia differentiation, and consequently, compromise the sense of smell (for example, see McEwen, Koenekoop et al. 2007, Bergboer, Wyatt et al. 2018, Uytingco, Green et al. 2019). Interestingly, therapeutic interventions to ameliorate ciliary defects in ciliary disorders currently holds the most promise for anosmia. Since the OE is exposed to the environment, the ease of its accessibility has been exploited for adenoviral-based gene delivery in clinically relevant mouse models, such as for intraflagellar transport (IFT)88 mutation or Bardet-Biedl syndrome (BBS), with encouraging results (McIntyre, Davis et al. 2012, Williams, Uytingco et al. 2017, Xie, Habif et al. 2021). Thus, these breakthroughs offer a tangible prospect of gene therapy for curing smell deficits in ciliopathy patients.

There are several attributes of the olfactory cilia that make them distinct from sensory or motile cilia in other cell types. The two distinctive morphological characteristics are a) they are considerably long and filamentous, reaching lengths of up to 200 μm in some species (Reese 1965), and b) they are produced in multiple numbers, about 10-30/cell in mice, arising from multiple basal bodies in the dendritic knob (McEwen, Jenkins et al. 2008, Jenkins, McEwen et al. 2009). In the mouse and the zebrafish, at the ultrastructural level, cilia from OSNs stand somewhere in between the conventional immotile primary cilia and the motile cilia: they have a (9+2) axonemal microtubule arrangement (like motile cilia), but lack the dynein arms that confer motility (like primary cilia) (Menco 1984, Ringers, Olstad et al. 2020, Pinto, Rasteiro et al. 2021).

As mentioned above, genes that have a generic function in ciliary differentiation, such as those encoding components of the key ciliary transport processes, IFT and the BBSome, are also required for proper formation of olfactory cilia (McIntyre, Davis et al. 2012, Williams, McIntyre et al. 2014). However, despite the importance and clinical significance of the olfactory cilia, how OSNs are transcriptionally programmed to generate this unique cilia-type has not been established. In the worm *C. elegans*, the RFX domain containing transcription factor Daf-19 is required for olfactory cilia biogenesis (Swoboda, Adler et al. 2000). Vertebrate genomes possess multiple paralogs of the Rfx family, and some of them have been linked to primary as well as motile cilia differentiation in specific organs and tissues (Choksi, Lauter et al. 2014). In addition, the vertebrates utilize the forkhead transcription factor Foxj1 for programming cells to generate motile cilia (Stubbs, Oishi et al. 2008, Yu, Ng et al. 2008, Choksi, Lauter et al. 2014). Whether these regulatory proteins are also involved in the generation of the specialized immotile olfactory cilia and contribute to their distinctive ultrastructure has thus far remained unexplored.

In this study, we show that in the zebrafish as well as the mouse, *Foxj1* genes are expressed in the differentiating OSNs, and animals with loss-of-function mutations exhibit a profound disruption in olfactory ciliogenesis. Through transcriptomics analysis as well as direct visualization of gene expression in the OSNs, we found that target genes regulated by Foxj1 in motile ciliated cells that encode components of the ciliary motility complex are repressed. This finding provides a molecular logic of how Foxj1 could be involved in differentiation of the immotile olfactory cilia. Rather intriguingly, besides affecting ciliary differentiation, loss of Foxj1 also affected the development of the OSNs themselves. Notably in the zebrafish, we were able to identify that expression of *ompb* (Danciger, Mettling et al. 1989, Çelik, Fuss et al. 2002, Sato, Miyasaka et al. 2005, Dibattista, Al Koborssy et al. 2021) and *cnga4* (Bradley, Reuter et al. 2001, Munger, Lane et al. 2001), encoding two key OSN-specific proteins involved in olfactory signal transduction, were strongly diminished in the absence of Foxj1 activity. In line with this, *foxj1* mutant zebrafish larvae showed reduced response to bile acid that is detected by the activity of Omp-positive OSNs (Friedrich and Korsching 1998, Sato, Miyasaka et al. 2005, Kermen, Franco et al. 2013, Bergboer, Wyatt et al. 2018). Thus, in addition to the previously established role of Foxj1 in motile ciliogenesis, we describe here novel functions for Foxj1 in regulating the differentiation of the immotile olfactory cilia and in controlling the development of the OSNs themselves, and consequently the OE.

## RESULTS

### Foxj1 is expressed in zebrafish OSNs

In our earlier analysis of the transcriptional pathways regulating the generation of the motile multiciliated cells (MCCs), we noted that one of the zebrafish orthologs of mammalian *Foxj1, foxj1b*, is expressed robustly in the developing OSNs (Chong, Zhang et al. 2018, D’Gama, Qiu et al. 2021). To examine this aspect in further detail, we used a transgenic gene trap strain, *Gt(foxj1b:GFP)*^*tsu10Gt*^, which contains GFP inserted within the first intron of *foxj1b* and faithfully reports the transcription pattern of the gene (Tian, Zhao et al. 2009). Consistent with our earlier observations using mRNA *in situ* hybridization, robust GFP expression could be detected in the differentiating OSNs of the larvae, co-labelled using an antibody against the neuronal RNA binding protein HuC (Figure 1A, B). Expression of *foxj1b* in OSNs is retained to adulthood, as shown by robust GFP expression throughout the lamella of the adult OE that contains the OSNs (Figure S1A, B) (Sato, Miyasaka et al. 2005). By contrast, we identified that the paralogous gene, *foxj1a*, is predominantly expressed in the motile cilia bearing MCCs that line the periphery of the larval olfactory cavity (Figure 1C) or the tips of the adult OE, and much less prominently in the OSNs (Figure S1B). For this analysis of *foxj1a* expression, we used a newly generated gene trap line (*Gt(foxj1a:2A-Tag-RFP)*) where Tag-RFP was inserted into the *foxj1a* locus and recapitulates faithfully the endogenous expression pattern of *foxj1a*. MCCs also expressed *foxj1b*, shown with the antibody against glutamylated-tubulin, one of the building blocks of motile cilia (Figure 1D).

**Figure 1:**
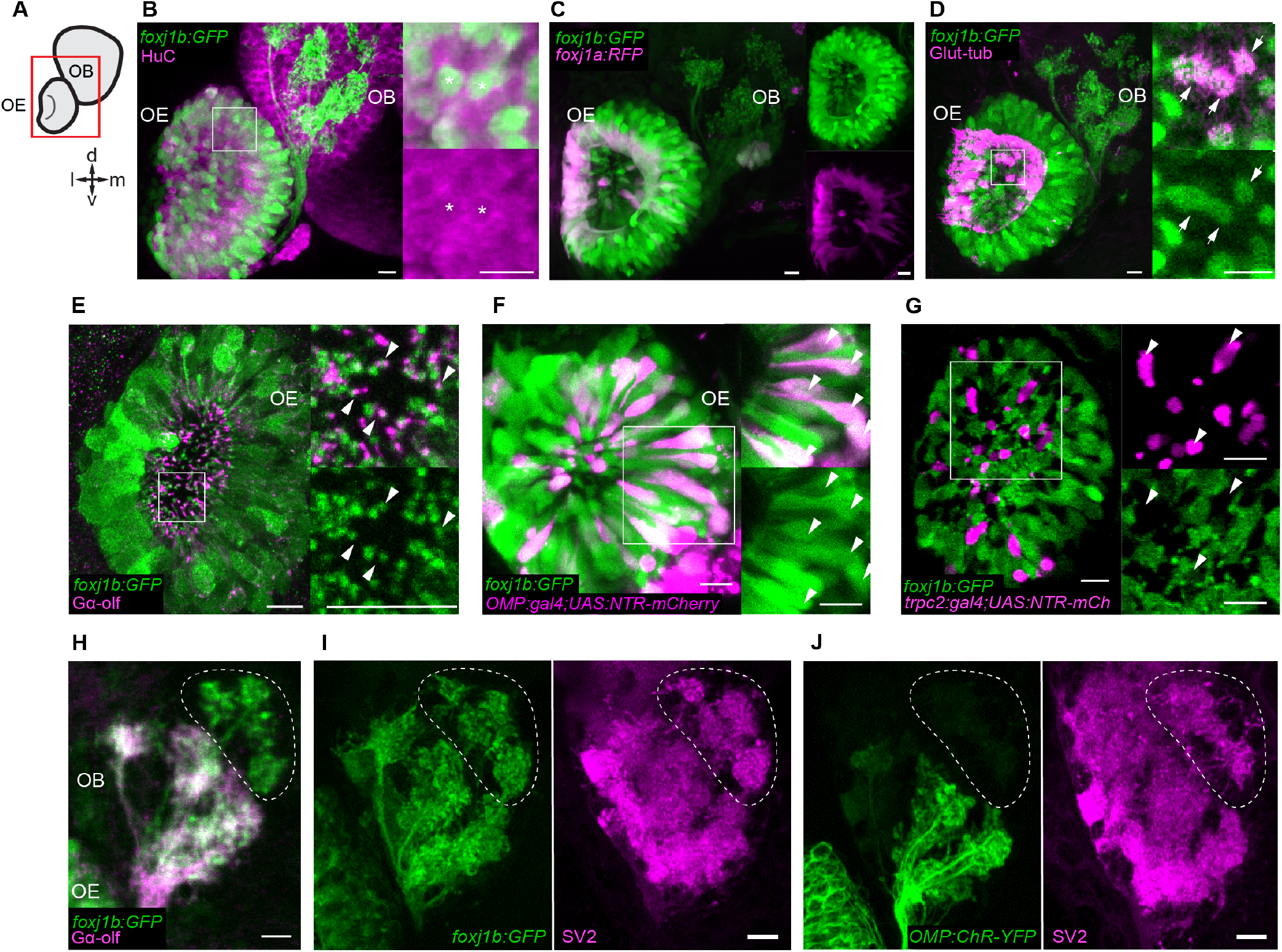
foxj1 is expressed in the zebrafish OSN. (A) Schematic showing the larval zebrafish olfactory epithelium (OE) and olfactory bulb (OB). (B-J) Confocal images of the larval (4-5dpf) zebrafish nose and olfactory bulb. (B) foxj1b (Gt(foxj1b:GFP), green) is expressed in neurons labelled by HuC (magenta). Asterisks show foxj1-expressing neurons. (C) Differential expression of foxj1a (Gt(foxj1a:2A-TagRFP), magenta) and foxj1b (Gt(foxj1b:GFP), green) paralogs. Note that foxj1a is expressed primarily at the outer rim of the olfactory cavity. (D) foxj1b-positive cells (Gt(foxj1b:GFP), green) bear brush of motile-cilia marked by glutamylated-tubulin (magenta). (E) foxj1b-positive OSNs (Gt(foxj1b:GFP), green) in the nasal pit bear cilia, indicated by the marker Gαolf (magenta). White arrowheads in insets show Gαolf marked cilia arising from foxj1b-positive OSNs. (F) Overlap of foxj1b-expressing cells (Gt(foxj1b:GFP), green) with the ciliated OSN marker OMP (Tg(OMP:gal4);Tg(UAS:NTR-mCherry), magenta) in 5dpf larva. (G) Mutually exclusive localization of foxj1b-expressing cells (Gt(foxj1b:GFP), green) and microvilli OSN, indicated by the marker trpc2 (Tg(trpc2:gal4);Tg(UAS:NTR-mCherry), magenta). (H) foxj1b-expressing OSNs (Gt(foxj1b:GFP), green) project mainly to Gαolf-labelled glomeruli (magenta). Note that foxj1b:GFP and Gαolf do not overlap entirely (dotted region). (I-J) foxj1b-positive OSNs (I) project to more glomeruli than OMP-expressing OSNs (J) (Tg(OMP:ChR-YFP)) (J). Glomeruli are shown by staining with the presynaptic vesicle marker SV2 (magenta). Scale bars = 10μm, l: lateral, m: medial, d: dorsal, v: ventral

Zebrafish OSNs are thought to largely comprise two subtypes – the ciliated OSNs and the microvillar cells, which are intermingled in the OE (Sato, Miyasaka et al. 2005). To determine the cell type specificity of *foxj1* expression, we examined foxj1b-GFP expression with markers of either ciliated (olfactory marker protein (Omp)) and the G protein Gαolf or microvillar OSNs (the channel protein Trpc2b).

Immunostaining with Gαolf revealed that the *foxj1b* expressing OSNs extend Gαolf positive-cilia into the nasal cavity (Figure 1E, note that only the base of the cilia are indicated in the projection to highlight the colocalization with GFP). We further confirmed that the GFP-positive cells in Gt(*foxj1b:GFP*) are ciliated OSNs by co-localizing GFP with mCherry expressed under the activity of *omp* regulatory elements using a combination of the *Tg(omp:gal4)* and *Tg(UAS:NTR-mCherry)* transgenes (Koide, Miyasaka et al. 2009, Agetsuma, Aizawa et al. 2010) (Figure 1F). In parallel, we investigated if the *foxj1b*-expressing OSNs could be microvillous OSNs using embryos transgenic for *trpc2b-gal4; UAS-Ntr-mCherry* (DeMaria, Berke et al. 2013). We found a mutually exclusive expression pattern of GFP and mCherry, implying that the *foxj1* genes are not expressed in the *trpc2b*-expressing microvillar OSNs, but primarily in *omp*-expressing ciliated OSNs (Figure 1G).

GFP expression also filled the afferent axons of the OSNs and could be traced right up to their synaptic arborizations on the glomeruli of the OBs. Interestingly, we did not observe a complete co-localization of GFP and Gαolf signals in the OB of larval zebrafish, suggesting that *foxj1b* is expressed in another OSN subtype (Figure 1H). To better decipher the identity of these additional *foxj1b* expressing OSNs, we stained the larval and adult OBs with the presynaptic marker, synaptic vesicle 2 (SV2), which specifically labels the glomeruli (Buckley and Kelly 1985, Braubach, Fine et al. 2012, Braubach, Miyasaka et al. 2013). We observed that *foxj1b*-expressing OSNs project to a broader range of glomeruli than OMP-expressing OSNs, including the ventral medial glomeruli (vmG) and ventral posterior glomeruli (vpG)(Figure 1I-J, Figure S1C-D) (Braubach and Croll 2021).

### Foxj1 is expressed in mouse OSNs

Similar to the zebrafish, we successfully identified Foxj1 expression within the mouse OE using an established antibody that specifically recognizes the Foxj1 protein of mammals (Pan, You et al. 2007) (Figure 2A). MCCs in the respiratory epithelium were strongly labelled for Foxj1 in accord with previous observations (Blatt, Yan et al. 1999). Within the OE, Foxj1 was detected in mature OSNs as demonstrated by overlap with the OSN-specific marker OMP (Figure 2A, C), albeit at a several fold lower expression level than in the MCCs (Figure 2D). The data aligned well with an earlier report of the abundant expression of *Foxj1* mRNA in adult mouse OE (Sammeta, Yu et al. 2007). Importantly, no Foxj1 immunostaining was detected in the OE of the *Foxj1* knockout mice confirming the specificity of the signal produced by the antibody (Figure 2B). Foxj1, being a transcription factor, appeared localized predominantly in the nuclei of the OSNs (Figure 2C, arrows; Figure S2). We also compared the relative level of Foxj1 expression in the OSNs to that observed in the MCCs and found no significant difference at different post-natal stages of development - P0, P5 and P30 (Figure 2D, Figure S2).

**Figure 2:**
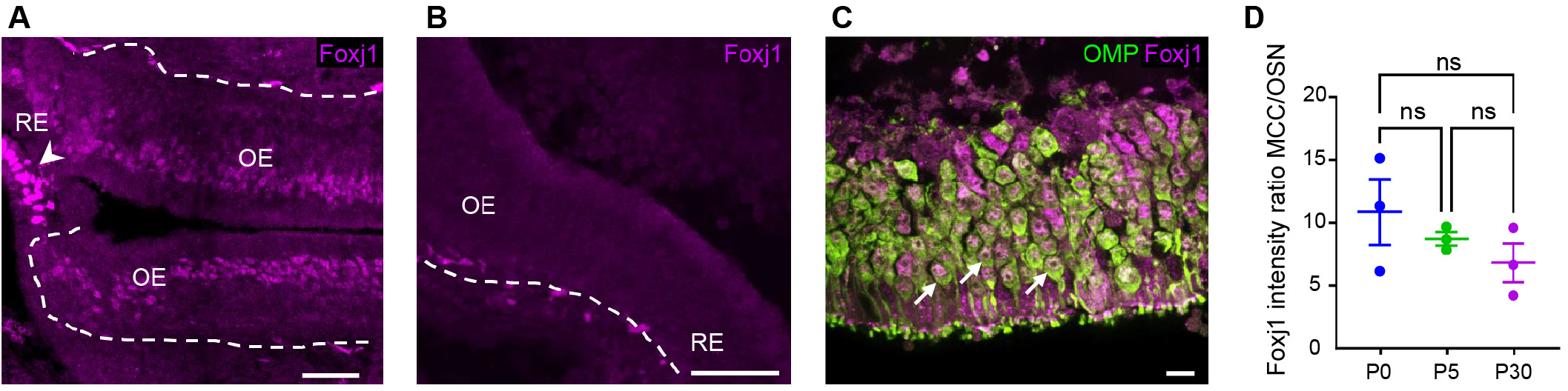
Foxj1 is expressed in the mammalian OSN. (A) In the nasal epithelium of adult mouse, Foxj1 localizes to multiciliated cells (arrowhead) within the respiratory epithelium (RE) and the layer of OSNs in the olfactory epithelium (OE). Dashed line demarcates lamina propria separating OE from the underlying tissue. (B) Foxj1 is absent in the nasal epithelium of the knockout Foxj1 mouse. (C) Foxj1 (magenta) localizes to the nuclei of mature OSNs immunostained for the olfactory marker protein (OMP). (D) Expression level of Foxj1 in OSNs at different age (newborn P0, day 5, P5 and adult P30) calculated by comparing immunofluorescence in OSNs relative to respiratory multiciliated cells (MCCs). No significant difference was found (MCC/OSN ratio of 10.87±2.61, P0; 8.74±0.53, P5; 6.83±1.56, P30, 3 mice in each group) by one-way ANOVA with Kruskal-Wallis test. Scale bars: 50 µm (A,B) and 10 µm (C).

### Foxj1 is required for olfactory cilia biogenesis in zebrafish

Having established that Foxj1 transcription factors are expressed in vertebrate OSNs, we next examined if they are required for olfactory ciliogenesis. Foxj1 has a central role in motile cilia biogenesis in diverse vertebrates including the zebrafish, *Xenopus*, mice, and humans (Stubbs, Oishi et al. 2008, Yu, Ng et al. 2008, Choksi, Lauter et al. 2014). Extrapolating from these earlier findings, it seemed reasonable to expect that *foxj1* expression in the OSNs could be linked to the generation of the olfactory cilia. As introduced above, the G-protein Gαolf is enriched on zebrafish and mammalian olfactory cilia and is a useful marker to label these organelles and distinguish them from the motile cilia of the neighbouring MCCs. Using Gαolf staining, we detected a reduction in the numbers of OSN cilia in *foxj1b*, but not *foxj1a*, homozygous larvae (Figure 3A-A”). However, when we analysed *foxj1a/b* double mutant larvae, we observed a profound loss of the olfactory cilia, indicating that redundancy between the two paralogs prevents the manifestation of a strong phenotype in the individual mutants (Figure 3A’’’). Despite the ciliary defects, we did not observe any aberration in the innervation of the Gαolf-positive OSNs to the OB in the double mutant larvae (Figure 3B-B’).

**Figure 3:**
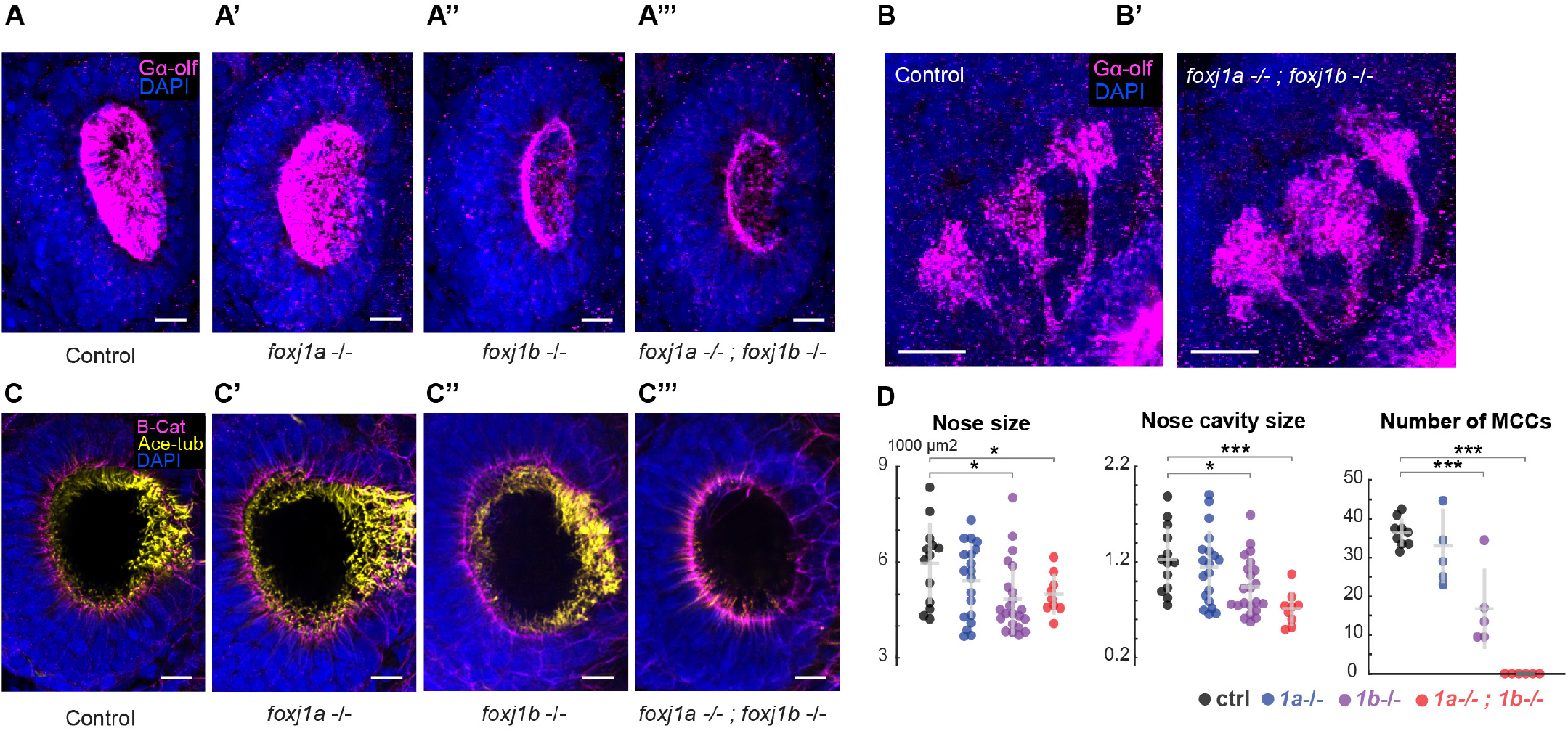
Foxj1 is required for olfactory cilia and motile cilia biogenesis. (A-A’’’) Immunostaining with anti-Golf antibody marking olfactory cilia (magenta) in foxj1a,b mutant larvae at 4 dpf. Nuclei marked with DAPI (blue). (A’) foxj1a mutants show no observable effect on the formation of olfactory cilia (n=3). (A”) foxj1b mutants show reduced olfactory cilia (n=3). (A”‘) foxj1a/b double mutants showing severe loss of olfactory cilia (n=3). (B-B’) Innervation pattern of ciliated OSN labelled with Gαolf (magenta) in control (n=3) and foxj1a/b double mutant (n=5) show no significant effect in the formation of olfactory glomeruli at 4 dpf. Nuclei marked with DAPI (blue). (C-C’’’) foxj1a/b double mutant showing severe reduction of motile cilia number as shown by immunostaining with anti-acetylated-tubulin for marking cilia (yellow) and beta-catenin for marking cell borders (magenta) in foxj1a,b mutant larvae at 4 dpf. (C’’’). Nuclei marked with DAPI (blue). (D) Significant decrease in the size of nose and nasal cavity, and number of MCC in foxj1b and foxj1a/b double mutant embryos. (For nose size, nControl = 13, nfoxj1a-/- = 20, nfoxj1b-/- = 22, nfoxj1a-/- ; foxj1b-/- = 9. For number of MCCs, nControl = 9, nfoxj1a-/- = 5, nfoxj1b-/- = 5, nfoxj1a-/- ; foxj1b-/- = 6). Scale bars = 10 μm.

In addition to affecting ciliary differentiation in the OSNs, motile cilia of the MCCs along the lateral edges surrounding the nasal pits (Reiten, Uslu et al. 2017, Ringers, Bialonski et al. 2023) were also completely absent in *foxj1a/b* double mutants, consistent with the established function of the Foxj1 proteins in motile cilia differentiation (Figure 3C-C’’’). We also identified that the loss of cilia in *foxj1a/b* double mutants coincided with a significantly reduced size of the nasal cavity and OE (Figure 3D), suggesting a possible impact of these mutations on the morphological structuring of the nasal cup or on OSN differentiation.

### Foxj1 is essential for establishing the OE in mice

Remarkably, loss of *Foxj1* in P21 mice resulted in a severe disfigurement of the olfactory system at the gross anatomical level, largely undeveloped turbinates and the entire nasal cavity was filled with DAPI-positive cellular masses composed of neutrophils (Figure S3A, A’, C). At the level of the OE, loss of *Foxj1* strongly impacted its cellular composition. In the mutants, the apical ciliary layer of the epithelium was completely disrupted and largely absent, in part, due to the significant loss of the mature OMP-positive OSNs themselves (Figure 4A-A’). Similar to the mature OSNs, the population of immature GAP43-positive OSNs was also disorganized and greatly depleted (Figure 4B-B’). Loss of both types of OSNs also resulted in a decreased thickness of the olfactory epithelium. Moreover, loss of *Foxj1* resulted in a substantial decrease of active proliferation in the olfactory epithelium as measured by Ki67-positive cells, compared to the wild-type (Figure 4C-C’). However, such a profound pathological remodeling of the olfactory epithelium in the Foxj1 knockout mice was not accompanied by cell death as reported by a cleaved Caspase (cCas9) immunofluorescence (Figures S3B-B’).

**Figure 4:**
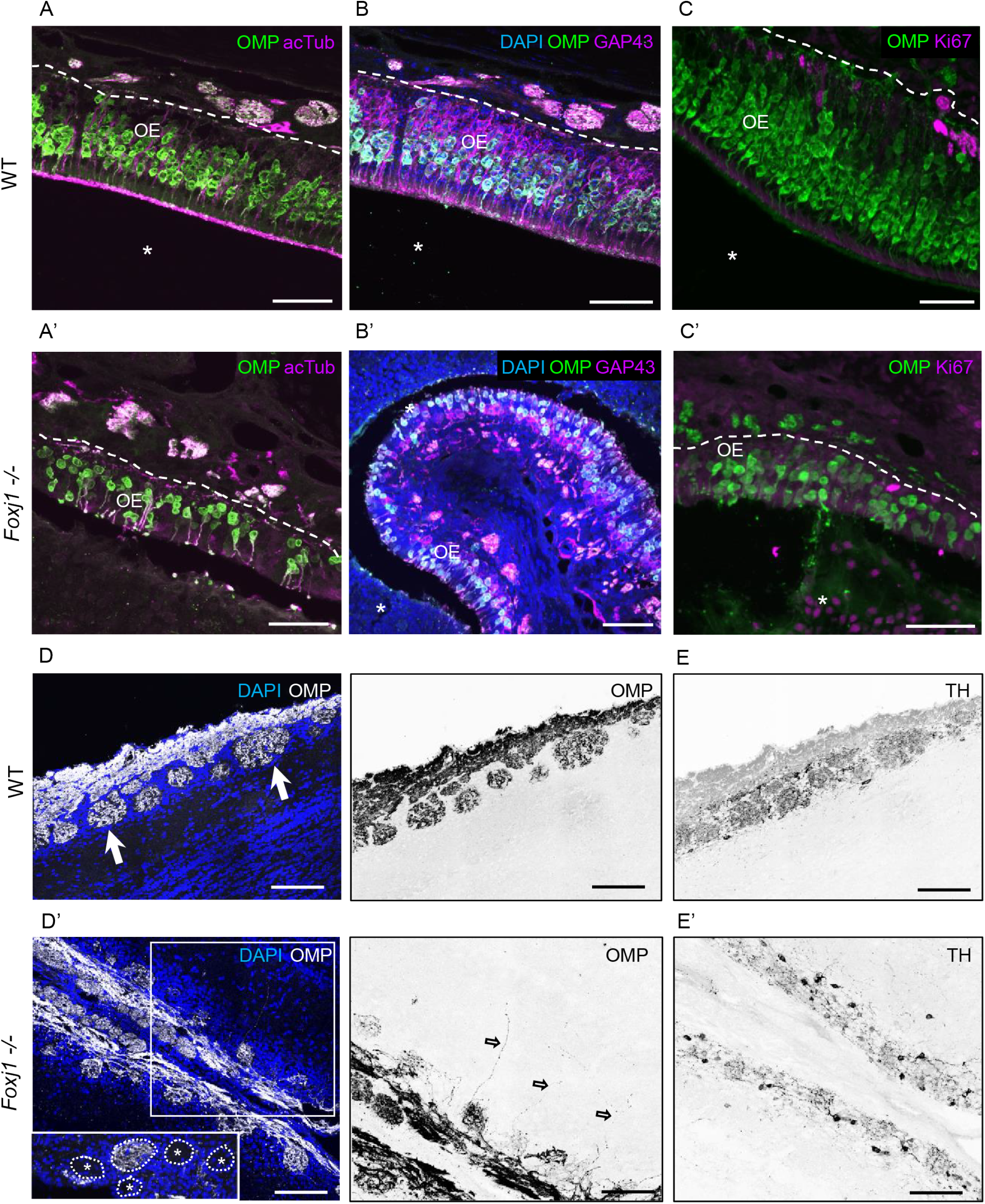
Foxj1 is essential for the establishment and ciliation of the olfactory epithelium in mice. (A-A’) Representative image of a control (A) and Foxj1-/- (A’) mouse OE stained for OMP (green), a mature OSN marker. Apical layer composed of mucus and cilia is strongly labelled for acetylated α-tubulin (magenta) in control (A). (A’) Mature OSNs (OMP, magenta) were fewer in number and disorganized within the OE of the Foxj1-/- (10,630±923 cells per mm2, n=8, WT; 3,106±714, n=7, KO, p=0.0006). Compared to the WT, thickness of the OE was significantly reduced in the Foxj1-/- (79.68±4.31 µm, n=34, 3 mice, WT; 45.48±2.15 µm, n=30, 3 mice, p<0.0001, KO). (B-B’) Immature OSNs (GAP43, magenta) were located below the layer of mature OSNs (OMP, green) as shown in a control mouse (B). (B’) Immature OSNs (GAP43, magenta) lost their orientation relative to mature OSNs and were fewer in numbers within the OE of the Foxj1-/- as compared to controls (B) (6,339±1,015 cells per mm2, n=8, WT; 2,008±481, n=7, KO, p=0.0012). The nasal cavity (A-C’, asterisks) was unobstructed in the WT whereas in the Foxj1-/- it was completely filled with DAPI and Ki67 positive cells. (C-C’) Proliferating cells expressing Ki67 (magenta) were mostly comprised of basal progenitor cells lining lamina propria. (C’) Fewer proliferating cells (Ki67, magenta) were present in the OE of the Foxj1-/- (C’) as compared to the control (C) (2,362±321 cells per mm2, n=9, WT; 502±72, n=9, KO, p<0.0001). (D-D’) Representative images of the olfactory bulb having oval shaped glomeruli (arrows, left) filled with the axonal projections of OSNs (OMP, white) in control (D). (D’) Reduced intensity of OMP immunostaining in the Foxj1-/- as compared to control (D) (352.5±26.7 a.u., n=21, WT; 266.6±17.5 a.u., n=44, KO; p = 0.0133). In the Foxj1-/- mouse glomeruli were less developed and had smaller perimeter than in the WT (2,558±368 µm, n=21, 2 mice, WT; 1,331±306 µm, n=44, 2 mice, KO; p<0.0001). Some glomeruli even contained no axonal projections as shown on a different representative image (D’ left, inset, asterisks). Axons did not fully converge within glomeruli overshooting in internal layers of the olfactory bulb (D’ right, arrows). (E-E’) Glomeruli were intensely labelled for tyrosine hydroxylase (TH), which reflects neuronal activity, in the WT mouse (E) as compared to Foxj1-/- (E’) showing weaker TH immunofluorescence (185.6±19.7 a.u., n=14, WT; 63.6±7.9 a.u., n=37, KO, p<0.0001). Scale bars = 50 µm (A-C), 100 µm (D-E’).

Given such a dramatic alteration in the structure of the OE, we asked if the loss of *Foxj1* affected the formation of the OB, which strongly depends on the strength of afferent innervation. Overall anatomy of the OB was not significantly affected in the *Foxj1* mutant mice (Jacquet, Salinas-Mondragon et al. 2009). However, it showed smaller sized glomeruli innervated by OSNs, at significantly smaller density than in the wild-type (Figure 4D, D’), as measured by the intensity of OMP immunofluorescence. Remarkably, some glomeruli completely lacked innervation by OMP-positive OSN axons, suggesting aberrant targeting by the OSNs (Figure 4D-D’, inset). Impairment of axonal targeting to the OB was further characterized by the existence of wandering axons unable to terminally synapse with glomeruli and stochastically projecting in the inner layers of the OB (Figure 4D’ right). This phenomenon is typical of young developing OBs and is not observed at P21 in the wild-type mouse (Wu, Ma et al. 2018). The aberrant innervation also resulted in a smaller size of glomeruli in the mutants. Furthermore, tyrosine hydroxylase (TH) immunostaining, a hallmark of normally functioning OB (Smith, Baker et al. 1991), was dramatically reduced in the knockout mice (Figure 4E-E’), suggesting a significant decrease of the overall neuronal network activity in the OB. Taken together, our results show that loss of Foxj1 in mice has a dramatic consequence on the differentiation of the OE and innervation patterns of the OBs.

### OSNs in zebrafish, mice and human do not express ciliary motility genes

Foxj1 transcriptional targets include a large set of genes that encode components for motile cilia biogenesis (such as members of the IFT machinery and dynein assembly factors), structural components of the motile ciliary axoneme (such as components of the radial spokes, dynein docking complex and central pair) and the axonemal dynein proteins themselves that confer motility (Choksi, Babu et al. 2014, Mukherjee, Roy et al. 2019). Since olfactory cilia are immotile and transmission electron microscopy (TEM) studies have also revealed that their axonemes lack dynein arms (Jafek 1983, Menco 1984, Hansen and Zeiske 1998, Ringers, Olstad et al. 2020), we next explored the mechanism by which such a variation to ciliary architecture is brought about, despite being controlled by Foxj1. When we analyzed for the expression of Foxj1 target genes in single cell RNA transcriptomes of OSNs obtained from the nasal cavities of zebrafish, mouse and human (Brann, Tsukahara et al. 2020, Durante, Kurtenbach et al. 2020, Kraus, Huertas et al. 2022), we found that there was a notable absence or severe reduction in expression of genes encoding axonemal dyneins and other associated motility components (Figures 5A and S4).

**Figure 5:**
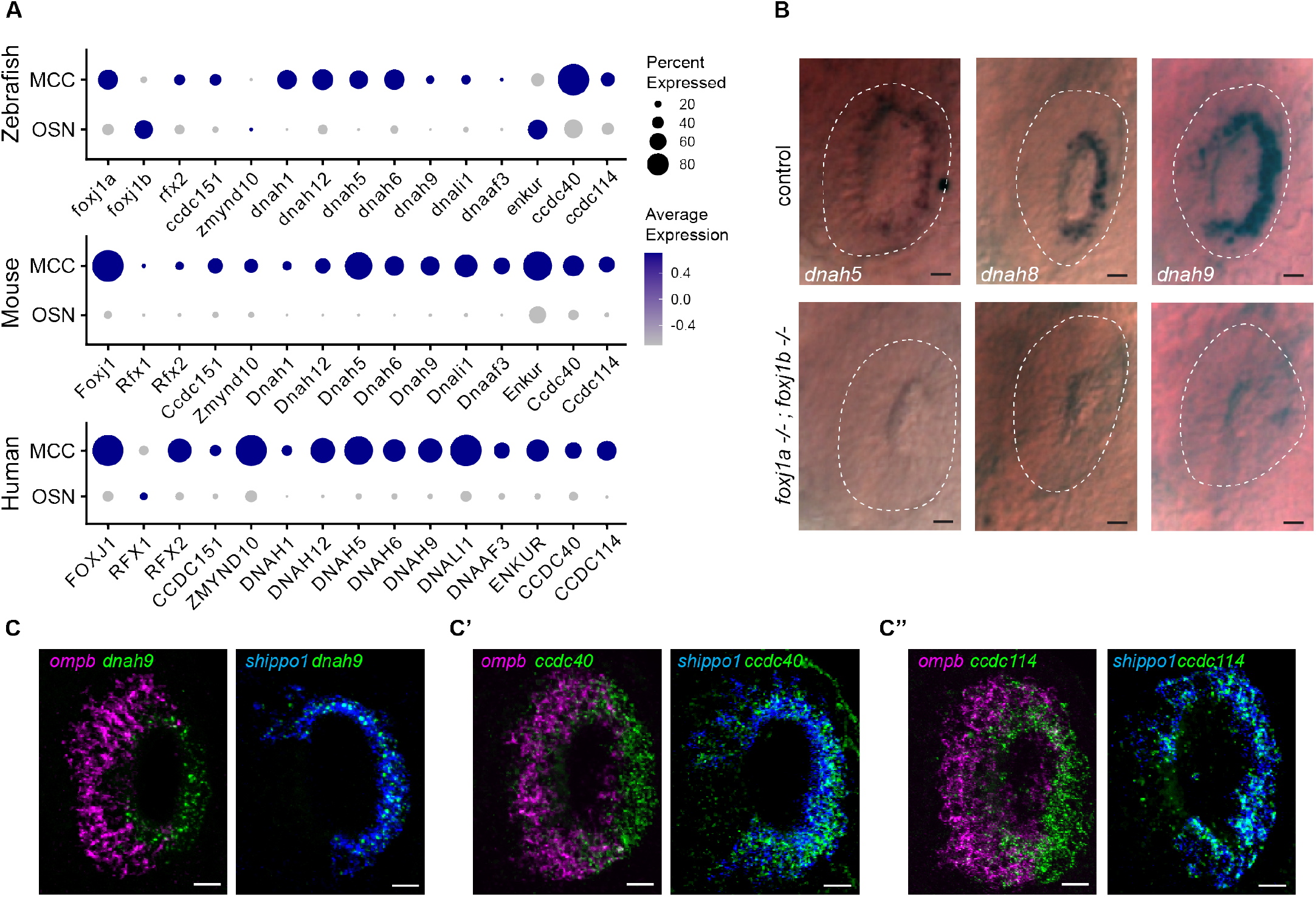
OSNs do not express axonemal dynein genes. (A) Single-cell transcriptome analysis of ciliated OSNs from zebrafish, mouse and human showing low expression of genes encoding axonemal dynein and motility-associated components in OSNs as compared to MCCs. (B) Whole-mount in-situ hybridization of dnah5, dnah8 and dnah9 in zebrafish embryo at 3 dpf showing absence of gene expression in the periphery of the nasal placodes (highlighted with white dashed line) that primarily consists of OSNs whereas expression is present in MCCs at the rim of the cavity (n=2 clutches). MCCs are depleted in foxj1a/b double mutants and therefore expressions of these genes were absent in the nasal placodes. Scale bars = 10μm. (C-C’’) Double fluorescent labelling of dnah9 (C), ccdc40 (C’) and ccdc114 (C’’) (green) with ciliated-OSN marker ompb (magenta) and MCC marker shippo1 (blue) by HCR in-situ hybridization in wildtype embryo at 72hpf showed co-expression of dnah9, ccdc40 and ccdc114 with shippo1 (n=3) but not with ompb (n=3). Scale bars = 20μm.

To experimentally validate these observations, we performed whole-mount in situ hybridization in zebrafish embryos for a collection of axonemal dyneins (*dnah5, 8, 9*), as well as *ccdc114* (encoding a dynein docking complex component) (Knowles, Leigh et al. 2013, Onoufriadis, Paff et al. 2013) and *ccdc40* (encoding a microtubule binding protein of motile cilia axonemes) (Becker-Heck, Zohn et al. 2011, Oda, Yanagisawa et al. 2014) to document their expression pattern in the MCCs versus the OSNs. Notably, we found that all of the dynein genes (*dnah5, 8, 9*) as well as *ccdc114* and *ccdc40* are expressed in the nasal MCCs (located at the rim of the nasal cavity and labelled with *shippo1*); however, in the OSNs (located throughout the nasal placode and labelled with *omp*), we failed to detect their expression (Figure 5B-C). Moreover, we further confirmed that in the MCCs axonemal dynein expression depends on Foxj1 activity (Figure 5B). Based on these observations we conclude that despite the necessity of Foxj1 in the olfactory ciliogenic program, there is a specific repression of its canonical motile ciliary gene targets in the OSNs.

### Foxj1 controls the expression of genes involved in OSN differentiation

To better understand the impact of loss of *foxj1* on OSN differentiation, we next performed RNA sequencing on 4 day old *foxj1a* and *foxj1b* mutant larvae. We observed that two OSN-specific markers, *ompb* and *cnga4*, were significantly downregulated in *foxj1b* but not in *foxj1a* mutants (Figure 6A, B). Omp is a small cytoplasmic protein thought to function in modulating odorant response in the OSNs (Danciger, Mettling et al. 1989, Çelik, Fuss et al. 2002, Sato, Miyasaka et al. 2005, Dibattista, Al Koborssy et al. 2021)), while Cnga4, a cyclic nucleotide gated subunit, plays a role in odorant-dependent adaptation (Munger, Lane et al. 2001). We further examined the status of expression of these genes using whole mount *in situ* hybridization in *foxj1a,b* double mutants. For both genes, we could confirm their dependence on Foxj1 activity: expression of both genes was almost completely lost from the double mutants (Figure 6C-D). Since *ompb* and *cnga4* encode important components of the olfactory signal transduction machinery, our observations underscore a previously unrecognized function of Foxj1 transcription factors in regulating aspects of the gene expression program specific for OSN differentiation.

**Figure 6:**
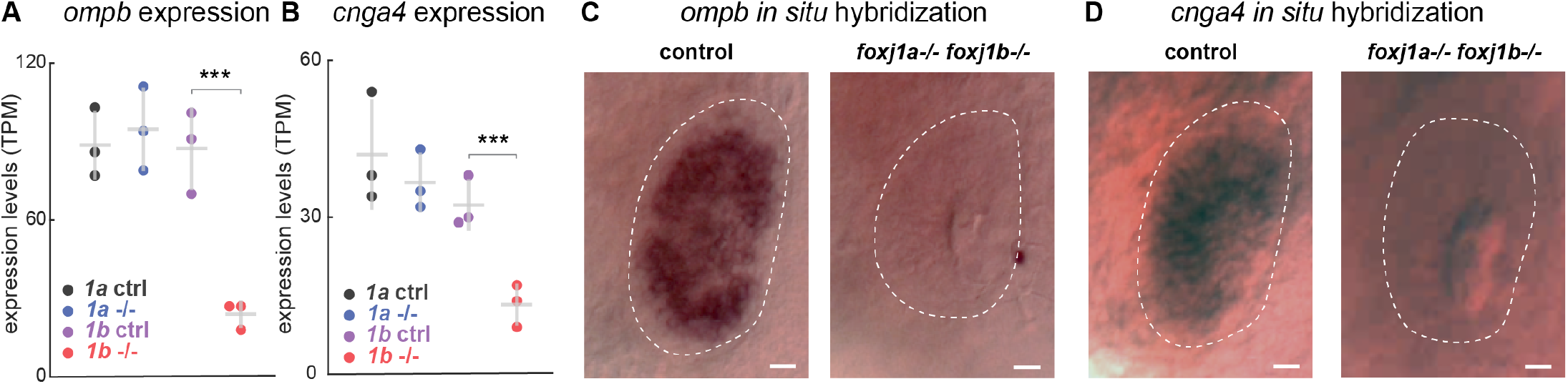
Foxj1 controls the expression of genes involved in OSN differentiation. (A, B) RNA sequencing of in foxj1a and foxj1b mutant embryos shows a significant decrease in the expression of the ciliated-OSN marker ompb (A) and cnga4 (B) in foxj1b mutant embryos but not in foxj1a mutants. (C, D) In-situ hybridization in 4 dpf larvae shows reduced expression of ompb (C) and cnga4 (D) in nasal placodes of foxj1a/b double mutant embryos. (n=3).

### Odor responses to bile acids are impaired in foxj1 mutant zebrafish larvae

As there was a dramatic change in OSN ciliogenesis in the *foxj1* mutants and we could, in addition, identify a role for Foxj1 in OSN differentiation beyond its requirement in ciliogenesis, we finally examined whether olfactory responses are impacted in the absence of its activity. For this, we subjected 4 day old zebrafish larvae expressing the calcium indicator GCaMP6s pan-neuronally (using the transgenic line *Tg(elavl3:gcamp6s))* to various odor types (Figure 7A, B). The odor set was composed of mixtures of amino acids, bile acid, nucleic acid and food odor that activate different subtypes of OSNs, and thus, their corresponding glomeruli (Friedrich and Korsching 1997, Friedrich and Korsching 1998, Sato, Miyasaka et al. 2005, Yaksi, von Saint Paul et al. 2009). Indeed, by investigating the response patterns in the OB, we were able to show responses to amino acids ventro-laterally, whereas bile acid-responding glomeruli were found medially, as previously described (Friedrich and Korsching 1997, Friedrich and Korsching 1998, Yaksi, von Saint Paul et al. 2009, Jetti, Vendrell-Llopis et al. 2014) (Figure 7C-C’). Since OSN responses to odor categories are highly selective and the various types of OSNs are intermingled in the epithelium (Figure 1F-G, 7C’), we measured the relative change in calcium signal in the entire OE as indicated in Figure 7B, rather than in individual cells. Using these assays, we showed that loss-of-function of *foxj1b* alone was not sufficient to impair responses of OSNs to any of the odor categories (Figure 7D-D’’’’). However, mutations in both *foxj1a* and *foxj1b* led to a significant difference in bile acid response (Figure 7E-E’’’’), implicating a compensatory role of each paralogue over the other. Since ciliated OSNs are known to be primarily responsible for detecting bile acids, whereas microvilli-containing OSNs detect amino acids and possibly nucleotides (Sato, Miyasaka et al. 2005, Bergboer, Wyatt et al. 2018), reduction in bile acid response is in line with our observations of impaired ciliogenesis and differentiation of ciliated OMP-positive OSNs.

**Figure 7:**
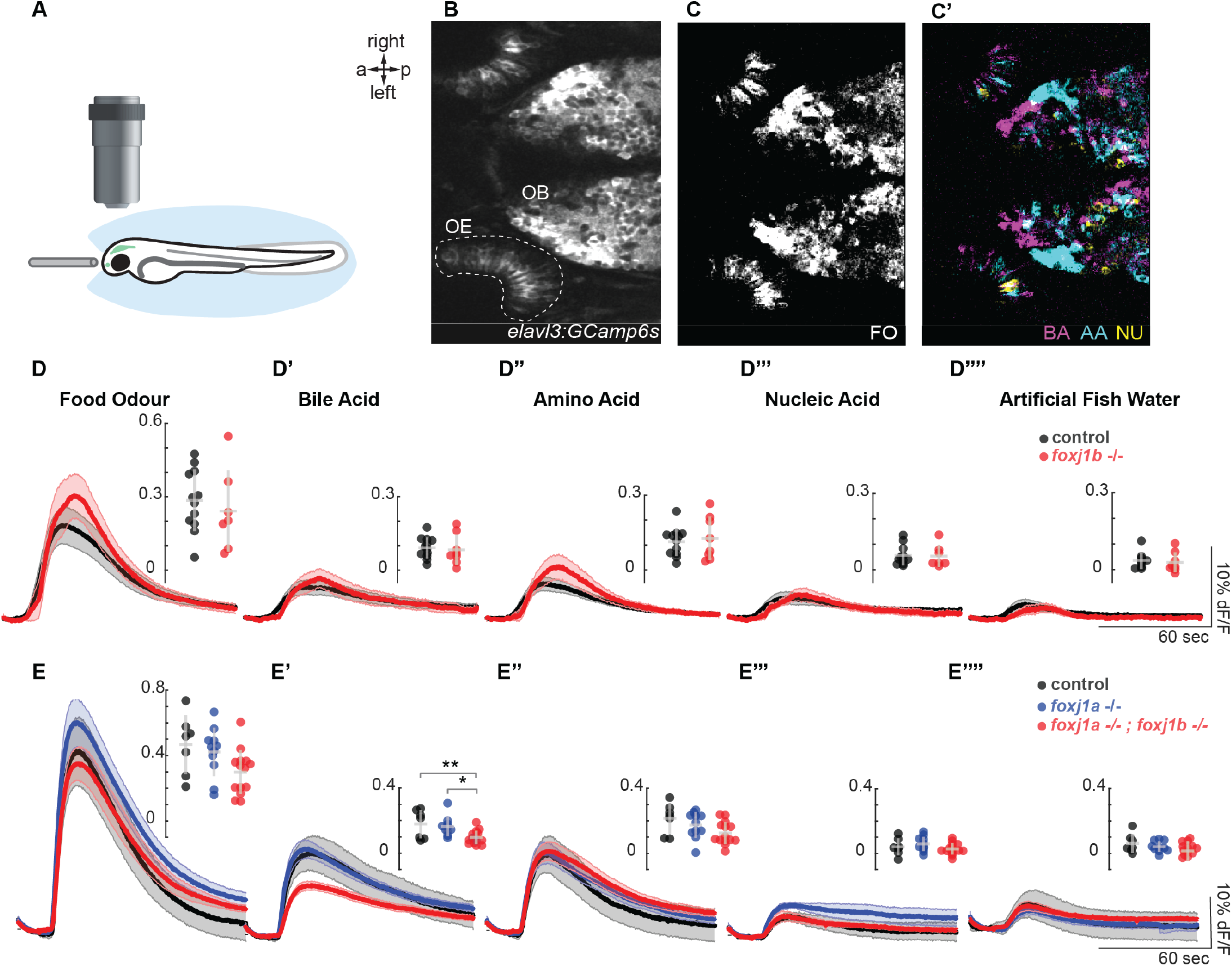
Odor responses to bile acid are reduced in foxj1 mutant zebrafish larvae. (A) Schematic of the olfactory experiment, showing a 4dpf zebrafish larva embedded in agarose. Its nose was exposed, and the odor stimuli were delivered by a fine tube. (B) Neural activity was measured using the Ca2+-reporter GCaMP6s (Tg(elavl3:Gcamp6s)) in a region of interest spanning the entire OE. (C-C’) Representative example showing neural activity in the OE and OB response to various odor types (FO: food odor, BA: bile acid, AA: amino acid, NU: nucleic acid). Note that responses in the OB are spatially organized, with non-overlapping domains. (D-D’’’’) Averaged traces of neural activity on the OE for foxj1b control (n=12) and mutant (n=8) fish for each odor type show no difference in odor responses. Maximum amplitude is shown in the insets. (E-E’’’’) Averaged traces of neural activity on the OE for foxj1 control (n=7), foxj1a mutant (n=10) and double foxj1a/b mutant (n=14) show a significant reduction in bile acid response. Maximum amplitude is shown in the insets. Significance identified by two-sample t-test, *: p<0.05, **: p<0.01. OE: olfactory epithelium, OB: olfactory bulb, a: anterior, p: posterior.

## Discussion

We perceive the world through our five senses of touch, smell, sight, taste and hearing. For many of these sensory modalities, cilia play a critical role. For example, photoreceptor neurons elaborate the connecting cilium to transport photopigments, whereas the kinocilium of sensory hair cells of the inner ear is required for proper polarization of the actin-based stereocilia that harbour mechanosensitive channels (Falk, Losl et al. 2015). Our sense of smell is also dependent on specialized cilia present on olfactory sensory neurons where odorant receptors are localized (Falk, Losl et al. 2015, McClintock, Khan et al. 2020). Likewise, motility functions of cilia, such as mucus clearance from the airways, circulation of cerebrospinal fluid and sperm propulsion also reflect the use of a wide assortment of cilia-types (Zhou and Roy 2015) . Understanding how such a great diversity of cilia-types is generated remains a key question in the field.

Ciliation is a rather late event in the development of the OSNs. After their specification, nascent OSNs transit through an immature stage that is characterized by the elaboration of the dendritic knob, generation of multiple basal bodies, migration and docking of these basal bodies with the apical membrane of the dendritic knob and finally, extension of the olfactory cilia (Ching and Stearns 2020, McClintock, Khan et al. 2020, Ching, Wang et al. 2022). Ciliation happens concurrent with the expression of genes necessary for the OSNs to mature into sensory neurons capable of transducing odor-evoked signals. (Nickell, Breheny et al. 2012, Heron, Stromberg et al. 2013, Hanchate, Kondoh et al. 2015). Thus, expression of transduction components like Omp and Cnga4 occur concurrently with the differentiation of the olfactory cilia. Anosmia is a prevalent symptom among many ciliopathy patients, and deficits in the ability to smell can also be inflicted by secondary insults to the olfactory cilia and the OSNs, such as upon viral infections (Uytingco, Green et al. 2019, Li, Li et al. 2020). Given these clinically relevant considerations, it is important to understand how olfactory cilia formation is developmentally regulated.

Using genetic analysis, we have now identified that the forkhead transcription factor Foxj1 is critical for OSNs to generate their specialized sensory cilia. We also found evidence that the expression of genes for ciliary motility proteins, such as the axonemal dyneins, is suppressed in the OSNs, clarifying why olfactory cilia are immotile and their lack of dynein arms in electron microscopic studies. In the zebrafish, the levels of *foxj1b* mRNA expression in the OSNs (judged by visual inspection and single cell RNA seq) is comparable to many cell types that differentiate motile cilia, negating an obvious “levels” issue and leading us to argue that the selective absence of ciliary motility gene expression in the OSNs likely arises from an active transcriptional repression mechanism. However, in the mouse, where we could directly analyse Foxj1 protein expression in the olfactory versus the respiratory epithelium, we observed that there is a stark difference, with several fold higher expression in the latter. Clearly, the molecular details underlying the transcriptional repression of cilia motility genes will need to be elucidated through future investigations. Of note, we have previously shown that *foxj1* is also required in auditory hair cells of the zebrafish ear for the differentiation of the kinocilia (Yu, Lau et al. 2011). Despite the name (kino = movement in Greek), kinocilia are not actively motile like the prototypical motile cilia. However, hair cells transcribe dynein genes and kinocilia have outer dynein arms but are believed to lack inner dynein arms (and hence the absence of active motility) (Wang and Zhou 2021, Shi, Beaulieu et al. 2023). Taken together, our identification of Foxj1 function in olfactory ciliogenesis further expands the role of this important transcription factor and suggests that its canonical motile cilia-specific transcriptional program is modified in different cells in different ways to generate a diversity of cilia-types that are suited for specific purposes. In this connection, it is also important to bear in mind that olfactory cilia in certain amphibian species have been described to be motile (Reese 1965, Ringers, Olstad et al. 2020) (see Supplementary video 1 for an example of a dissociated OSN from *Rana pipiens* (Kleene and Gesteland, 1991)). We speculate that in such instances, Foxj1 likely institutes the characteristic motile ciliogenic transcriptional program in the OSNs to facilitate the formation of motile olfactory cilia. Moreover, while cilia loss has only a mild, yet significant, impact in bile acid responses in the zebrafish as shown in our current and in an earlier study (Bergboer, Wyatt et al. 2018), loss of cilia function in mouse or human OSNs was associated with more severe anosmia (Kulaga, Leitch et al. 2004, McEwen, Koenekoop et al. 2007, Jenkins, McEwen et al. 2009). Taken together, this suggests that even though common principles dictate olfaction across vertebrates, there are some levels of species differences and sensitivity to cilia loss, that may relate to the ecological niche and need of individual species.

Interestingly, we have also found a more pervasive requirement of Foxj1 in OSN differentiation in the zebrafish as compared to the mouse, that transcends its traditional role in regulating ciliogenesis. In the zebrafish, genes for key olfactory signal transduction proteins, such as *ompb* and *cnga4*, which are expressed during the terminal differentiation process of the OSNs to become functional sensory neurons, are induced by Foxj1. This unexpected finding suggests that Foxj1 has been recruited to the OSN differentiation program independent of its role in regulating ciliary biogenesis. What additional OSN maturation genes are regulated by Foxj1 and whether this function is also relevant in the context of mammals remains to be clarified. In the mouse, in addition to ciliary defects, we observed that loss of Foxj1 had a dramatic consequence on the organization of the OE and innervation patterns of the OBs. In the zebrafish, however, we only observed minor changes in the size of the OE and no major innervation defects. It is important to note that we

## Materials and Methods

### Mouse strains

All procedures performed on mice were approved by the Washington University Institutional Animal Care and Use Committee (IACUC), protocol 22-0117. Knockout Foxj1 and wild-type littermates were housed in a standard animal facility. Wild-type mice of C57/B6 strain were obtained from the University of Florida mice breeding facility and handled according to the approved IACUC protocol 201908162.

### Zebrafish maintenance and strains

The animal facilities and maintenance of the zebrafish, *Danio rerio*, were approved by the NFSA (Norwegian Food Safety Authority) and the Singapore National Advisory Committee on Laboratory Animal Research. All the procedures were performed on zebrafish larvae of different developmental stages post fertilization in accordance with the European Communities Council Directive, the Norwegian Food Safety Authorities and the Singapore National Advisory Committee on Laboratory Animal Research. Embryonic, larval and adult zebrafish were reared according to standard procedures of husbandry at 28.5°C, unless mentioned otherwise. For our experiments, the following fish lines were used: *T2BGSZ10 Gt(foxj1b:GFP)* (Tian, Zhao et al. 2009), *Tg(OMP:Gal4;UAS:NTR-mCherry)* (Koide, Miyasaka et al. 2009, Agetsuma, Aizawa et al. 2010), *Tg(UAS:GCaMP6s)* (Muto, Lal et al. 2017), *Tg(OMP:ChR2-YFP)* (Sato, Miyasaka et al. 2005), could only investigate the innervation pattern and size of the OB at larvae stages due to the lethality of the *foxj1a/b* double mutants. Hence, it is still possible that *foxj1* regulates OB development later in development. The difference between zebrafish and mice may also relate to the early expression of Foxj1 during OB neurogenesis in mice (Jacquet, Muthusamy et al. 2011), but not in the zebrafish. Alternatively, it is possible that motile cilia-mediated flow, which is also impaired in the *Foxj1* mutant animals, may have a more significant impact on the morphogenesis of the OB in mice than the zebrafish. Further dissection of *Foxj1* function in the OSNs of the mouse will require selective ablation of the gene specifically from these neurons using conditional approaches.

## Acknowledgements

We would like to thank S. Kleene, University of Cincinnati, for the recording of ciliary beating on the dissociated frog OSN shown in Video S1, Y. Yoshihara for sharing the Gαolf antibody, and the fish facility support staff in our respective institutes for zebrafish maintenance. This work was supported by a FRIPRO research grant from The Research Council of Norway (NJY 314189) to N. J-Y. and the Agency for Science Technology and Research (A*STAR) of Singapore to S.R.

## Author contributions

Conceptualization: SR, NJY; Formal Analysis: DR, ME, KU, CR, YZ, SL, MIC; Funding acquisition: SR, NJY; Investigation: DR, ME, KU, CR, YZ, IY, PD, SL; Resources: SLB, HCP; Supervision: SR, NJY, SLB, JRM, EY, HCP, CK; Visualization: DR, ME, KU; Writing – original draft: SR, NJY, ME; Writing – review and editing: all authors.

*Tg(trpc2b:gal4)* (DeMaria, Berke et al. 2013), *Tg(UAS: NTR-mCherry)* (Davison, Akitake et al. 2007) *foxj1a* mutants (two different alleles generated in two different laboratories were used (Olstad, Ringers et al. 2019, Zhang, Jia et al. 2018)), *foxj1b* mutant (D’Gama, Qiu et al. 2021), *Gt(foxj1a:2A-TagRFP)*^*FRZCC 1100*^ (see below) and *Tg(elavl3:GCamp6s)* (Vladimirov, Mu et al. 2014).

Animals were analysed irrespective of their gender. Note that for zebrafish younger than 2-3 months, gender is not apparent. For adult stage imaging experiments, fish were selected according to their body size, which is reported in the manuscript, to ensure reproducibility of the results. Mutants were obtained either from heterozygous incross, heterozygous crossed with homozygous, or homozygous incross. We did not observe an impact of the parents’ genotype on the phenotype of the progeny. Controls were siblings with a control genotype (heterozygous or wild-type). For homozygous incross, controls were the progeny of a cross of wild-type siblings of the homozygous parents. Animals were genotyped prior to the experiments and their genotype was re-confirmed following the experiments. Animals were either in the AB or pigmentless *mitfa-/-* (Lister, Robertson et al. 1999) background.

### Generation of *Gt(foxj1a:2A-TagRFP)*^*FRZCC 1100*^ knock-in zebrafish

To generate the transgenic line to label *foxj1a*-expressing cells specifically, the short homology mediated knock-in strategy was utilized, as previously described (Wierson, Welker et al. 2020, Welker, Wierson et al. 2021). First, the Cas9 target site in exon 1 of *foxj1a* gene for the single guide RNA (sgRNA) was selected using CHOPCHOP (http://chopchop.cbu.uib.no/). The sgRNA was synthesized by the cloning-free gRNA *in vitro* synthesis method (Wierson, Welker et al. 2020, Jeong, Yun et al. 2021, Welker, Wierson et al. 2021), and Cas9 mRNA from the pT3TS-nCas9n vector (Addgene plasmid #46757) with mMESSAGE mMACHINE™ T3 transcription kit (Thermo Fisher). To validate the efficiency of the sgRNA, *foxj1a* sgRNA and Cas9 mRNA was injected into one-cell embryos and genomic DNA was extracted from the injected embryos at 24 hpf. Then, the T7 endonuclease I assay was performed as described in the prior reports (Jao, Wente et al. 2013, Jeong, Yun et al. 2021). Next, the homology arms were designed with the GTagHD (http://www.genesculpt.org/gtaghd/) and the pGTag-TagRFP-SV40 plasmid (Addgene) was used as a donor vector. The *foxj1a* targeting donor vector containing 2A-TagRFP-SV40 (Wierson, Welker et al. 2020, Welker, Wierson et al. 2021) was constructed and the plasmid was purified with the QIAGEN Plasmid Midi kit (QIAGEN) and PureYield Plasmid Miniprep System (Promega). 2 nl of a mixture of *foxj1a* sgRNA (50 pg), universal gRNA (25 pg), Cas9 mRNA (200 pg) and the target donor plasmid (10 pg) was microinjected into one-cell stage embryos. Then, the embryos which expressed red fluorescence in the *foxj1a*-expressing cells were screened and raised to adulthood (F0). To identify germline-transmitted lines, the F0 fishes were crossed with wild-type and whether the F1 embryos expressed *foxj1a*-specific red fluorescence or not was validated. After the germline-transmitted founder was screened by fluorescence, we confirmed how the DNA is integrated into the genome with the genomic DNAs from the larvae. We verified that the 2A-TagRFP-SV40 DNA is properly integrated into the target site by performing PCR at the 5’ and 3’ junctions of integration sites and sequencing.

### Two-photon calcium imaging

For *in vivo* imaging, fish were paralysed through α-bungarotoxin injection (Reiten, Uslu et al. 2017, Diaz Verdugo, Myren-Svelstad et al. 2019) and were embedded in 0.75% low-melting-point agarose (LMP, Fisher Scientific) in a recording chamber (Fluorodish, World Precision Instruments). To ensure odor delivery to the nostrils, the LMP agarose in front of the nose was carefully removed, after letting solidification for 30 min.

A two-photon microscope were used for calcium imaging: Scientifica, with a Nikon 16x NA 0.8 water immersion objective. For excitation, a mode-locked Ti:Sapphire laser (MaiTai Spectra-Physics) was tuned to 920nm (Diaz Verdugo, Myren-Svelstad et al. 2019). Recordings were performed as single plane recordings. The acquisition rate was 31Hz with an image size of 512x510 pixels. Data analysis was done with MATLAB as described in the subsequent section.

### Odor preparation

Our odor selection consisted of food odor, bile acid mixture (taurocholic acid, taurodeoxycholic acid), amino acid mixture (Alanine, Phenylalanine, Aspartic acid, Arginine, Methionine, Asparagine, Histidine), nucleic acid mixture (Inosine monophosphate, Adenosine monophosphate). All odorants were purchased from Sigma Aldrich. Food odor was prepared using commercially available fish food; 1g of food particles was incubated in 50ml of artificial fish water (AFW) for at least 1 h, filtered through filter paper, and diluted to 1:50. Amino acid mixture, bile acid mixture and nucleotide mixture were all prepared from the frozen stocks the day before use at 0.1mM and were kept at 4°C. All odor mixtures were allowed to equilibriate to room temperature before the experiments.

### Odor delivery

Odors were delivered with a tube positioned right in front of the nose for 20s through a constant flow of AFW (2ml/min). The stimulation was performed with HPLC injection valve (Valco Instruments) controlled with Arduino Due (Reiten, Uslu et al. 2017). Before each experiment, a trial with fluorescein (10-4M in AFW) was performed to determine precise onset of odor delivery. The four odor mixtures, as well as AFW were presented in a randomized order and this order of delivery was repeated thrice in total per fish. After the experiment, the larvae were retrieved for health-check and genotyping.

### Data analysis

Once the neural activity was recorded, the data were aligned. Distinct ROIs corresponding to the left and right olfactory epithelia were drawn. The change of fluorescence was estimated as the relative change of fluorescence over time by dF/F, F-F_baseline_/F_baseline_. F_baseline_ and was the average value of the frames at the onset of the recording before the stimulus delivery. A baseline of 10 seconds, ∼310 frames was used. The maximum of the response amplitude of the change of fluorescence was identified for each trial using the findpeaks function in the first minute of the recording and the average maximum was calculated across the left and right for the 3-trial repetitions.

Data analysis and statistics was done using MATLAB. Two-sample t-test was used for analysis. P<0.05 was considered as statistically significant.

### Immunostaining and confocal imaging

Immunostaining and H&E staining of the mouse olfactory system

Adult mice used in this study were anesthetized with Ketamine/Xylazine and cardiac perfused using ice-cold PBS, pH7.4, immediately followed by ice-cold 4% PFA in PBS. Air from nasal cavity was removed by vacuum to enhance access of the fixative to the OE. Neonate mice at P0 and P5 were anesthetized on ice, decapitated and heads drop-fixed in ice-cold fixative. The tissue was post-fixed overnight, decalcified in 0.5 M EDTA, pH8, incubated in 10%, 20% and 30% sucrose and frozen in Tissue-Tek OCT embedding medium (Fisher Scientific, cat# NC1862249). Importantly, immunodetection of Foxj1 required fixation in 1% PFA. Fixed tissue was sectioned at 12 µm thickness and sections were mounted on a Superfrost glass slides (Fisher Scientific Cat# 12-550-15). Except for Foxj1, immunostaining of all other marker proteins did not require an antigen retrieval. However, Foxj1 detection in mouse OE required heat-activated retrieval using Tris/EDTA (1mM EDTA, 0.05% Tween 20, pH 8.0) and incubation for 20 min at high pressure in a consumer grade pressure cooker (Instant Pot Duo). For immunostaining, tissue sections were washed 3x5min in PBS, then permeabilized for 15 min in PBS with 0.1% Triton X-100, and finally, incubated overnight at 4°C in blocking solution containing 2% normal donkey serum (Jackson Immunoresearch, cat# 017-000-121, RRID: AB_2337258), 0.5% bovine serum albumin (Sigma cat# A7030) prepared in PBS and 0.1% Triton X-100. Afterwards, tissue sections were incubated overnight at 4°C with primary antibodies diluted in the blocking solution. Dilutions of primary antibodies were as follows: anti-Foxj1 (1:100), anti-OMP (1:2,000), anti-acetylated tubulin (1:1,000), anti-Gap43 (1:500), anti-cleaved Caspase3 (1:500) and anti-TH (1:500). Next, tissues were washed 3x5min with PBS and incubated for 1h at room temperature with respective secondary antibodies (1:1,000 dilution). Finally, the tissues were counter-stained with DAPI and embedded in Prolonged Gold (Thermo Fisher cat# P10144). The processed tissues were analysed with an inverted confocal microscope Nikon TiE-PFS-A1R using pre-set optical configuration for DAPI, FITC, TRITC and Cy5. Images were post-processed using Nikon Elements software (version 5.11) and NIH ImageJ/FIJI.

Chromogenic H&E staining of the tissue was done according to the manufacturer’s instructions (Vector Laboratories H-3502).

### Analysis of immunofluorescence and morphology of the mouse OE

Number of cells immunostained with a respective marker protein was counted in several representative areas of the OE and then normalized to the area of the OE delineated by the basal lamina and uppermost apical layer. This approach accounted for the difference in the OE thickness in the wild-type and *Foxj1*-KO. Intensity of immunostaining for Foxj1 was measured in individual cells identified in the RE and OE, then grouped per type to compute an average and derive the ratio between respiratory MCCs and olfactory OSNs per each representative tissue area. Special care was taken to acquire immunofluorescence images of the wild-type and *Foxj1*-KO tissue at identical optical settings. Glomeruli in the OB were analyzed by measuring its perimeter whereby immunostaining was measured as an average fluorescence within each glomerulus delineated by DAPI staining. Statistical analysis of the data was done by Prism 9 (GraphPad). Unpaired two-tail Mann-Whitney test was used to compare two groups and p<0.05 was considered statistically significant. Three group comparison was done using one-way ANOVA with Kruskal-Wallis test.

### Immunostaining of larval zebrafish nose

4 dpf larval fish were fixed with a 4% paraformaldehyde solution (PFA) and 0.5% TritonX-100 in phosphate-buffered saline (PBSTx) at 4°C overnight. The samples were washed with 0.5% PBSTx to remove the fixing solution. 100% acetone was used for permeabilization of the larvae at -20°C for 20 mins, after which the samples were washed with 0.5% PBSTx (3x10min) and blocked with 0.1% bovine serum albumin (BSA) in 0.5% PBSTx at room temperature for 2h. Afterwards the samples were incubated with acetylated tubulin (1:1000, 6-11B-1, Sigma-Aldrich), glutamylated tubulin (1:400, GT335, Adipogen) or beta-catenin (1:200, Cell Signaling Technologies), HuC/D (1:150, A-21271, Invitrogen) antibodies overnight at 4°C. The next day, the larvae were washed (0.5%PBSTx, 3x1h) and subsequently incubated with the secondary antibodies (Alexa-labelled GAR555 plus and GAM IgG2b633, 1:1000, Thermo Fisher Scientific) in fresh blocking solution overnight at 4°C. On the third day, DAPI staining (1:1000, Thermo Fisher Scientific) was performed for 2h at room temperature in 0.5% PBSTx. The larvae were thoroughly washed (3x1h) in 0.5% PBSTx, then transferred in glycerol of increasing concentrations (25%, 50%, 75%) before being mounted in 75% glycerol. Confocal imaging was performed with Zeiss LSM 880 Axio Examiner Z1 with a Zeiss 20X plan NA 0.8 objective. For a detailed protocol, see D’Gama and Jurisch-Yaksi, 2023.

The same fixing, staining and imaging procedure was used on 5 dpf *foxj1b:GFP* larvae. These samples were incubated with the glutamylated tubulin (1:400, GT335, Adipogen) and anti-GFP-488 (1:1000, Thermo Fischer) antibodies and Alexa-labelled 555 plus (1:500, Thermo Fisher Scientific) secondary antibody.

### Immunostaining of the adult OB

For staining the OB of *foxj1b:GFP* fish, brain samples were dissected and fixed using 4%PFA in PBS and incubated at 4°C overnight. The next day, the samples were washed with 0.25%PBSTx (3x10mins). The samples were then incubated with 0.05% Trypsin-EDTA on ice for 40 min. After incubation, they were quickly washed twice and then for 10 min with 0.25%PBSTx. The samples were blocked with 2% DMSO+1% BSA made in 0.25% PBSTx for 4 hours. Samples were then incubated with mouse anti-SV2 monoclonal antibody (1:1000, DSHB), 1 ml per tube and agitated for 3-5 days at 4°C. The next day, the samples were washed with 0.25% PBSTx (3x1h). The samples were then incubated with secondary antibody diluted in 0.25% PBSTx, Alexa-labelled GAM555 plus (1:500, Thermo Fisher Scientific) + anti-GFP-488 (1:500, Thermo Fisher Scientific) + DAPI (1:1000, Thermo Fisher Scientific). This was incubated for 3 days at 4°C. The stained samples were washed with 0.25% PBSTx (3x1h) and were subsequently treated with increasing concentrations of glycerol (25%, 50% for 1h each). The stained samples were stored at 4°C and imaged using a Zeiss Examiner Z1 confocal microscope with a Zeiss 20x plan NA 0.8 objective.

### In-vivo confocal imaging of larval zebrafish

The in-vivo imaging of *foxj1a:RFP; foxj1b:GFP* larvae were performed at 5dpf. The larvae were immobilized using tricaine methanesulfonate (MS222) and were subsequently embedded in 1.5% low-melting-point agarose (LMP, Fisher Scientific), with the nose pointing up. After the agarose was solidified, confocal imaging was performed with Zeiss LSM 880 Axio Examiner Z1 with a Zeiss 20X plan water objective, NA 1. *In-vivo* imaging of f*oxj1b:GFP; OMP:gal4; UAS:NTR-mCherry* larvae was done similarly at 4 dpf.

### Quantification of nose morphology

Confocal images of 4 dpf larval noses were investigated with Fiji/ImageJ and their nose and nasal cavity size were measured. MCCs were counted making use of the acetylated tubulin and beta catenin immunostaining. All quantification was done on both nostrils of each animal and the average value across nostrils per larvae was reported. Statistical analyses were done using MATLAB. The two-sample t-test was used for analysis. P<0.05 was considered as statistically significant.

### Single-cell RNA sequencing analysis

Three publicly available scRNAseq datasets of the OE of different organisms: zebrafish (Kraus, Huertas et al. 2022), mouse (Brann, Tsukahara et al. 2020) and human (Durante, Kurtenbach et al. 2020) were selected and analyzed with Seurat (3.2.0) (Satija, Farrell et al. 2015, Butler, Hoffman et al. 2018, Stuart, Butler et al. 2019, Hao, Hao et al. 2021). The datasets consisted of 4561, 39985 and 27272 cells, respectively. Each dataset was clustered to identify main cell types using UMAP dimensionality reduction. Dims = 30 and resolution = 0.8 were used to form the clusters. For zebrafish, 21 cell clusters were obtained as a result. The following markers were used to identify main cell clusters: Neuron (*elavl3*), OMP-positive cells (*ompb*), Trpc2b-positive cells (*trpc2b*), Sustentacular cells (*krt5, lamc2, col1a1b*), Neural progenitors (*mki67*), Immune cells (*mpeg1*.*1, srgn*), MCCs (*dnah5l*), Supporting cells (*cfd, ccl25a, itga6b*), Blood cells (*hbba1*). Clusters 17, 18 and 19 could not be identified.

The Foxj1 target genes were identified as reported in our previous work (Choksi, Babu et al. 2014). Expression levels of selected Foxj1 target genes were identified in the cell clusters and were compared in the OSN and MCC clusters.

### RNA sequencing

To isolate RNA for sequencing, 4 dpf larvae were collected in a 1.5 ml tube and placed on ice. To lyse the samples, 500 μl trizol was added and the euthanized larvae were homogenized through a 27-gauge needle until the mixture looked uniform. After adding another 500 μl trizol, the samples were incubated for 5 mins at room temperature. The larvae were then treated with 200 μl chloroform, and the tube was rocked for 15 secs to mix the contents. The tubes were incubated for 2 mins at room temperature and then centrifuged for 15 mins at 12000 rpm at a temperature of 4°C. After centrifugation, the upper aqueous phase containing RNA was mixed with equal amounts of 100% ethanol and was then loaded onto an RNA spin column (Qiagen) and centrifuged for 30 secs at 8000 rpm. The spin column was further incubated with 700 μl of RW1 buffer and centrifuged for 30 secs at 8000 rpm. The spin column tubes were then placed into a new collection tube and further treated to remove any DNA contamination by washing the tubes with 350 μl of RW1 buffer followed by DNase enzyme (Qiagen) in RDD buffer (10 μl DNase+ 70 μl RDD buffer per tube) for 45 mins at room temperature. After incubation, 350 μl of RW1 buffer was added to the tubes and centrifuged for 15 secs at 8000 rpm. The tubes were then treated with 500 μl RPE buffer and centrifuged for 30 secs. This step was repeated twice, and the tubes were then centrifuged for 1 min at 8000 rpm to remove any residual buffer left in the column. For RNA extraction from the column, 30 μl nuclease free water was added and incubated for 2 mins. The tubes were then centrifuged for 1 min at 8000 rpm to elute the RNA. The concentration of the extracted RNA was quantified using Nanodrop and the quality was analysed with a bioanalyzer. The samples were then sequenced by BGI’s DNBSEQTM Technology. In order to analyze the RNA-sequencing data, the paired-end sequencing reads were aligned to the Zebrafish genome (GRCz11) using GSNAP (Wu and Nacu 2010) and read counts per gene were determined by using featureCounts (v2.0.3; (Liao, Smyth et al. 2014)) using ensemble version 106. Only paired-end reads that were uniquely mapped were kept for downstream analyses. The raw read counts were used as input for DESeq2 (v1.36.0, (Love, Huber et al. 2014)) for differentially expressed genes. Genes with abs(log2 fold change) > 2 and p-adj < 0.1 were assumed as significantly expressed genes. Datasets will be uploaded on GEO upon acceptance of the manuscript.

### Hybridization Chain Reaction (HCR)

HCR stainings were done according to the protocol by Choi et al. (Choi, Calvert et al. 2016) with some small modifications. All HCR probes, pre-hybridization buffer, amplification buffer, short-hairpins, wash buffers were purchased from Molecular Instruments, Inc. 15-20 zebrafish embryos at required stages were collected in 1.5ml eppendorf tubes and then fixed with 1 ml of 4% paraformaldehyde (PFA) for overnight at 4oC. Next day, embryos were washed for 2 hours with 1x phosphate buffer saline (PBS) with buffer change every half an hour. Finally, the embryos were transferred in absolute methanol and then stored in -20oC for overnight incubation. Next day, embryos were serially rehydrated by washing with 1 ml of 75%, 50% and 25% methanol diluted in PBS and subsequently with PBST (1x PBS with 0.1% Tween-20) washes few times for 5 minutes. For each wash, tubes were placed on a horizontal shaker. After the washes, embryos were treated with 1 ml of Proteinase K prepared in PBT (0.8 µl of 25 mg/ml Proteinase K in 1ml PBT). Treatment time varied according to the stages of the embryos. After the treatment, embryos were quickly washed for two times with 1 ml of PBST and finally residual Proteinase K activity was stopped completely by adding 1ml of 4% PFA. Embryos were incubated for 20 minutes at room temperature. Embryos were then washed three times with PBST for 5, 15 and 25 minutes, respectively. After the final wash, embryos were incubated in 350 µl of HCR pre-hybridization buffer (pre-hybe) pre-warmed at 45oC by placing the tubes in a floater and incubating in a water bath set at 45oC for 2 hours. After the incubation, 250 µl of probe solution (prepared by adding 1 µl of two or more HCR probes each tagged with different initiators, in 250 µl of pre-warmed pre-hybe; final concentration of 2 pmol of probe) was added in each tube. Embryos were incubated in the same water bath overnight (12-16 hours).

Next day, extensive washing was done. Embryos were first washed with 500 µl of HCR probe wash buffer for 2 times x 10 minutes each and thereafter 2 times for 30 minutes each with 70%, 50% and 25% wash buffer (diluted in 5x SSCT) and final one 15 min wash with 5x SSCT. All washes done by incubating embryos in 45oC water bath. Final two washes of 10 minutes done with 5x SSCT at RT. Embryos were then incubated for an hour in 500 µl HCR amplification buffer at RT. This buffer has been prewarmed at RT once taken out from 4oC storage. Meantime, HCR hairpin RNA solution was prepared by snap-cooling the hairpins. 10 µl of each h1 and h2 hairpins (both 30 pmol) for a single initiator was aliquoted in separate PCR tubes. Tubes were then placed in a heat block at 95oC for 1.5 minutes and immediately cooled at RT and incubated for 30 minutes in dark. Both h1 and h2 were then aliquoted in 250 µl of amplification buffer and mixed well. Once embryo incubation in amplification buffer was over, hairpin mixed buffer was then added to each tube and incubated at room temperature overnight (12-16 hours) in the dark.

Next day, amplification buffer mix was removed, and embryos were washed 5 times for 30 minutes with 1ml of 5x SSCT at RT. This was then followed by another 4 washes with 1 ml PBST buffer for 25 minutes each. 0.3 µl of DAPI (5 mg/ml) was added in the first wash with PBST. Once PBST washes were over, embryos were finally kept in 1 ml of 70% glycerol prepared in PBS and stored in -20oC. Embryos were imaged using an Olympus FLUOVIEW FV3000 upright confocal microscope.

### Whole mount in-situ hybridization (WISH)

WISH was done according to the routine protocol. Antisense-RNA probes for *ompb* was synthesized from *ompb* CDS cloned in the TOPO-TA vector while cnga4 probe was synthesized according to the protocol by Hua et al., 2018 (Hua, Yu et al. 2018). Primer sequences for *ompb* and *cnga4* probe synthesis are mentioned in the oligonucleotide section.

### DNA isolation for zebrafish embryo genotyping

Antibody-stained or FISH-stained embryo genotyping was done in two steps – DNA isolation and PCR followed by agarose gel electrophoresis. Genomic DNA isolation from individual embryos or parts of embryos such as excised head or torso was done by alkaline lysis method. A single embryo or embryo head or torso was transferred to individual PCR tube containing 20 µl of PBS. PBS was then removed and 20 µl of 25 mmol of alkaline lysis solution (25 mmol of sodium hydroxide + 0.2 mmol EDTA, prepared in sterile water) was added in the tube. The tube was transferred to a thermocycler for incubation at 95oC for 45 minutes. After incubation, the tube was taken out, vortexed and quick spun in a table-top spinner. 20 µl of neutralization buffer (40 mmol of Tris-HCl prepared in sterile water, pH 8.0) was then added to the tube and mixed well. The DNA mixture was now ready for PCR. 1 µl of this DNA mix was used for a 10 µl of PCR reaction.

For genotyping of larval zebrafish used for the nose quantification and odor-response experiments, the samples were subjected to gDNA isolation using 50 μl PCR lysis buffer (containing 10 mM tris pH7.5, 50 mM EDTA, 0.2% Triton X-100 and 0.1mg/ml Proteinase K) overnight at 50°C. To stop the reaction the samples were heated to 95°C for 10 minutes. The samples were then centrifuged at 13000 rpm for 2 mins. The supernatant containing gDNA was used for qPCR. For performing qPCR, 10 μl SYBR green PCR master mix (Thermofisher) was mixed with 0.5 μl each of forward and reverse primer and 7 μl water to make a 18 μl reaction mixture. This mixture was added to a 96 well qPCR plate (Thermofisher) and 2 μl of extracted gDNA were mixed with this reaction. The samples were then analyzed based on their melting curves as wild-type, heterozygous or homozygous or by further gel electrophoresis.

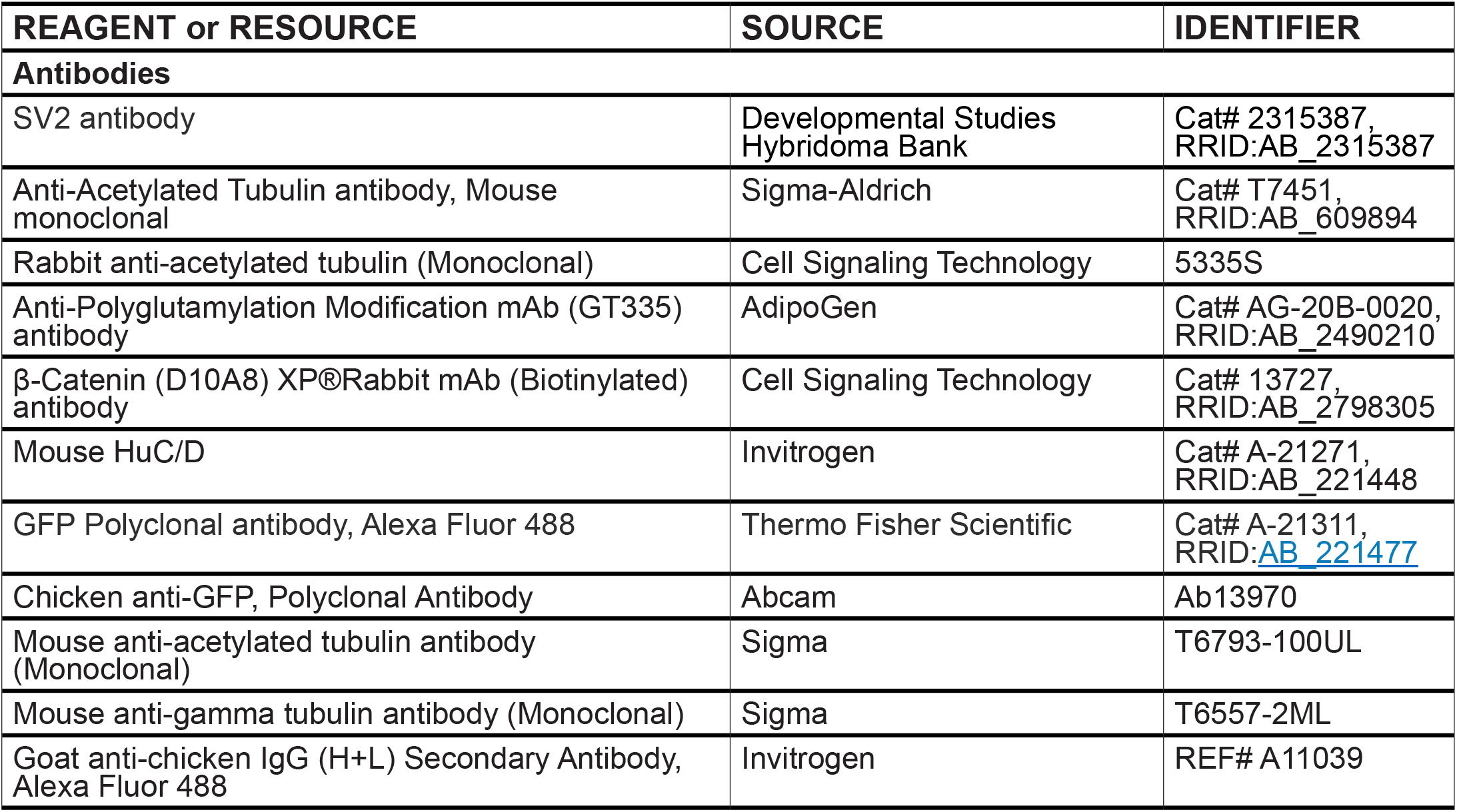
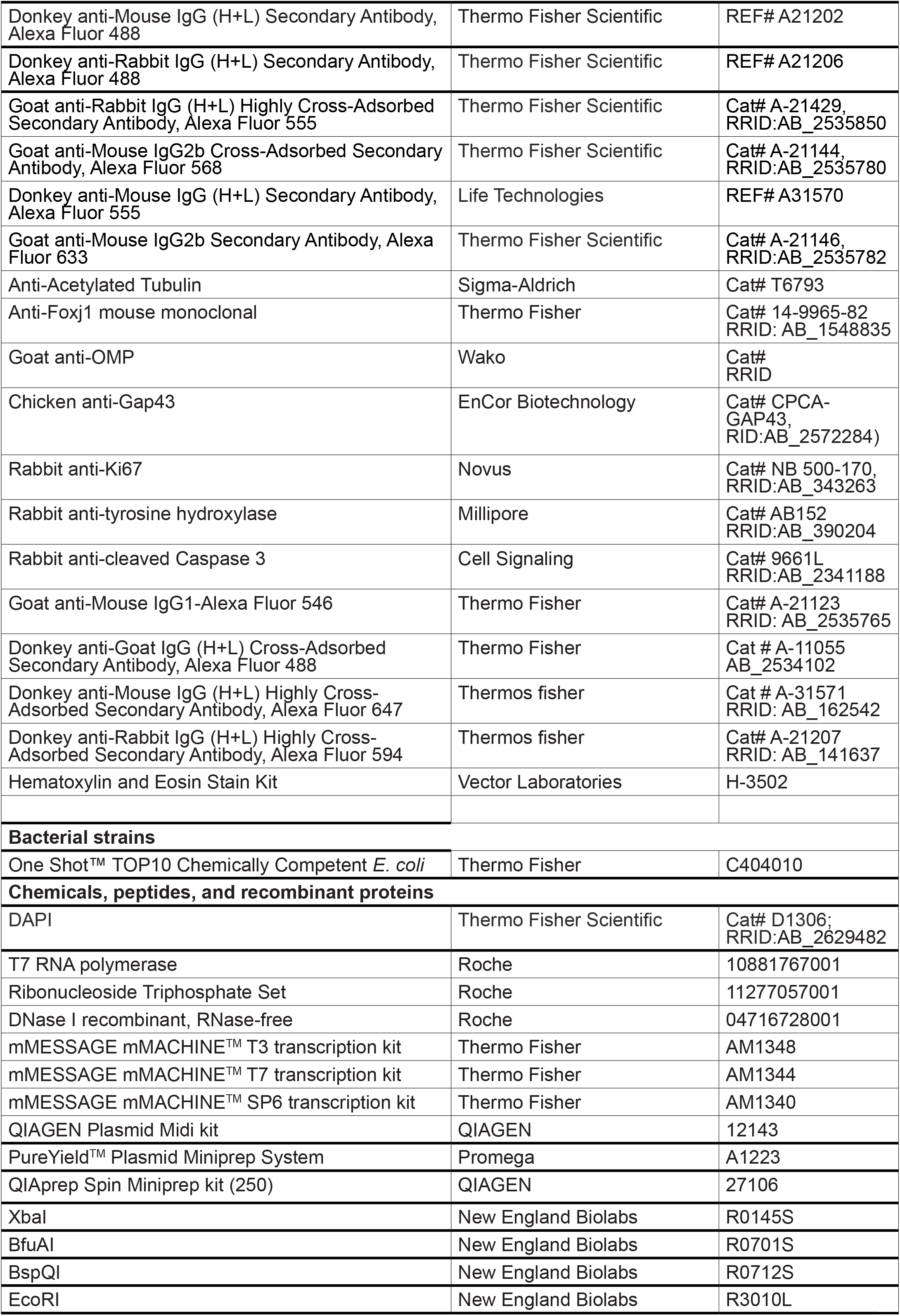
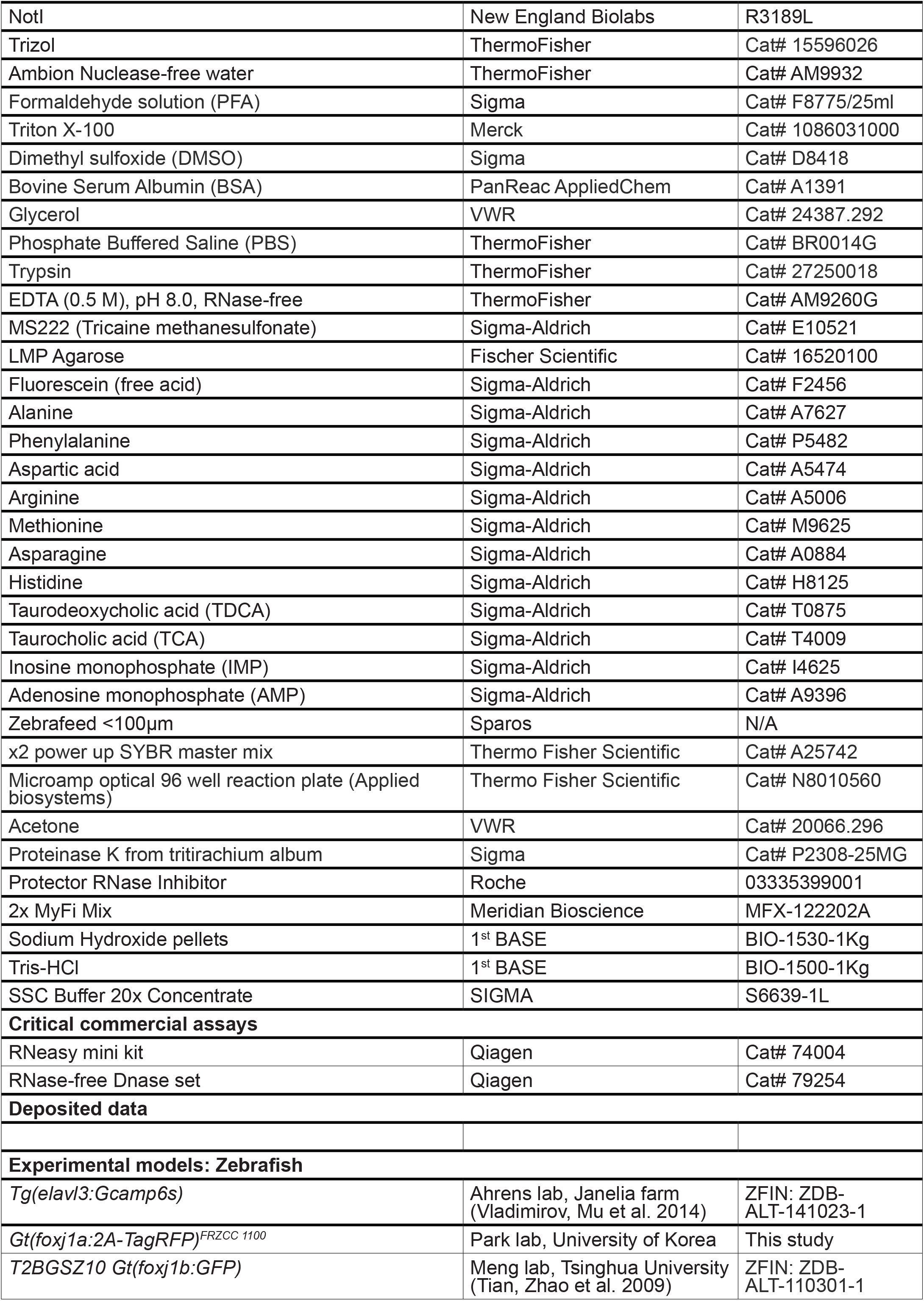
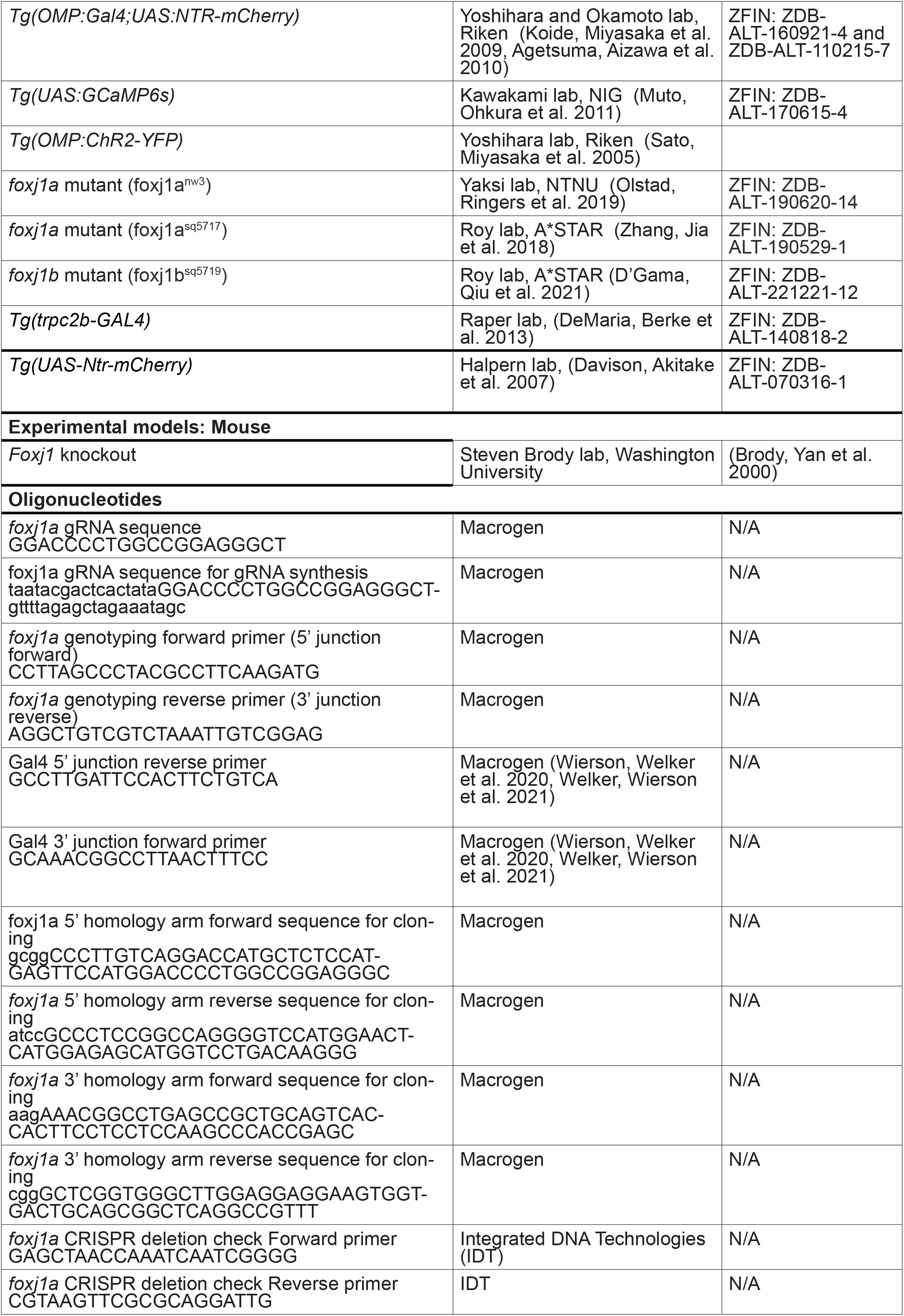
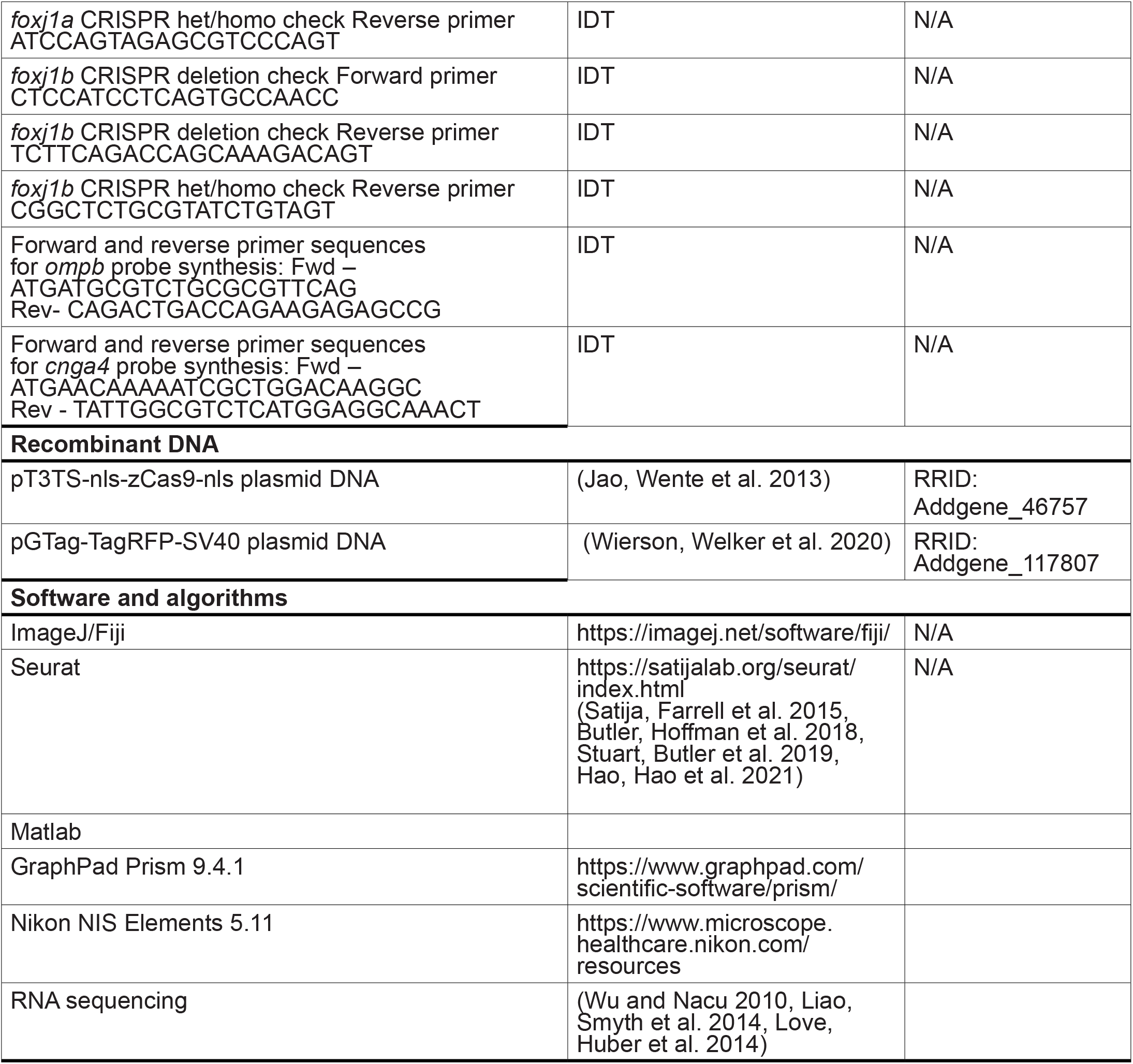

