## Supplemental Figures for "Foxj1 controls olfactory ciliogenesis and differentiation program of the olfactory sensory neurons"

**Figure S1: *foxj1* expression in the olfactory epithelium and projection of *foxj1*-expressing cells into the OB in adult zebrafish.**

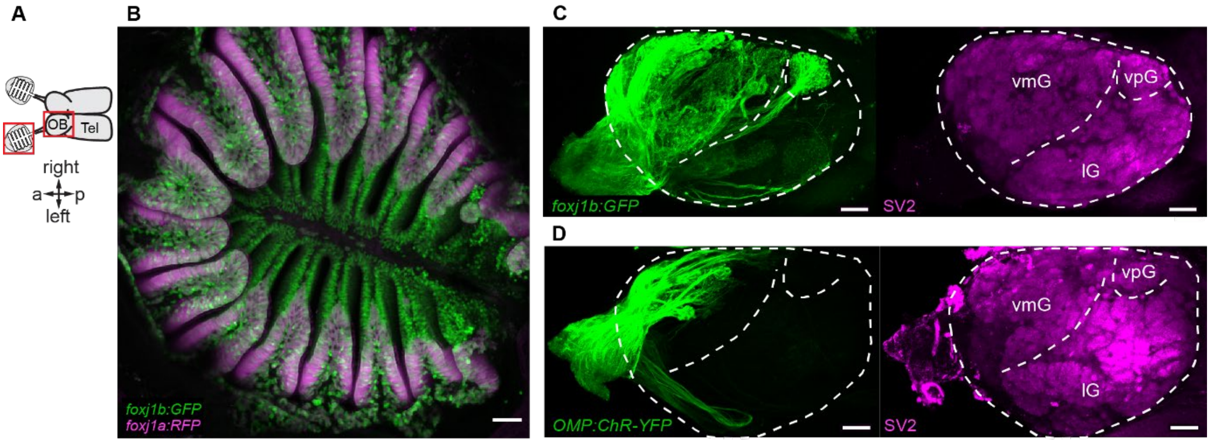


(A) Schematic showing the adult zebrafish olfactory bulb (OB) connected to the olfactory epithelia.
(B) Confocal image of an adult zebrafish olfactory epithelium (OE) shows expression of *foxj1a* (*Gt(foxj1a:2A-TagRFP)*, magenta) and *foxj1b* (*Gt(foxj1b:GFP)*, green) in OE. Note that *foxj1a* in mainly expressed at the tip of the lamellas where MCC are located. Scale bar = 50µm.
(C-D) Projections of *foxj1b*- (C, *Gt(foxj1b:GFP)*, green) and *omp*-positive OSNs (D, *Tg(OMP:ChR-YFP)* green) into the OB. Glomeruli are indicated by the presynaptic marker SV2. Note that *foxj1b*-expressing OSNs project to more glomeruli than *omp*-expressing OSNs. Scale-bars = 20µm. a: anterior, p: posterior

**Figure S2: Foxj1 expression in the olfactory epithelium at different age.**

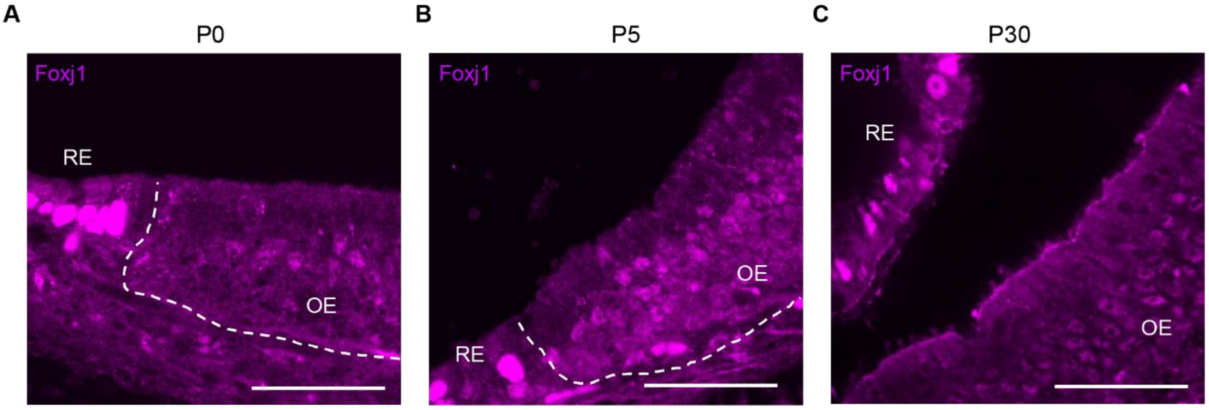


(A-C) Immunostaining of Foxj1 at different animal age (newborn P0 (A), day 5, P5 (B) and adult P30 (C)) in OSNs of wild-type mice. Border line between the olfactory epithelium (OE) and respiratory epithelium (RE) is marked by a dashed line. Brightly labelled cells in the RE are multiciliated respiratory cells. Scale bar = 50 µm.

**Figure S3: Loss of Foxj1 results in neutrophils infiltration in the nasal cavity**

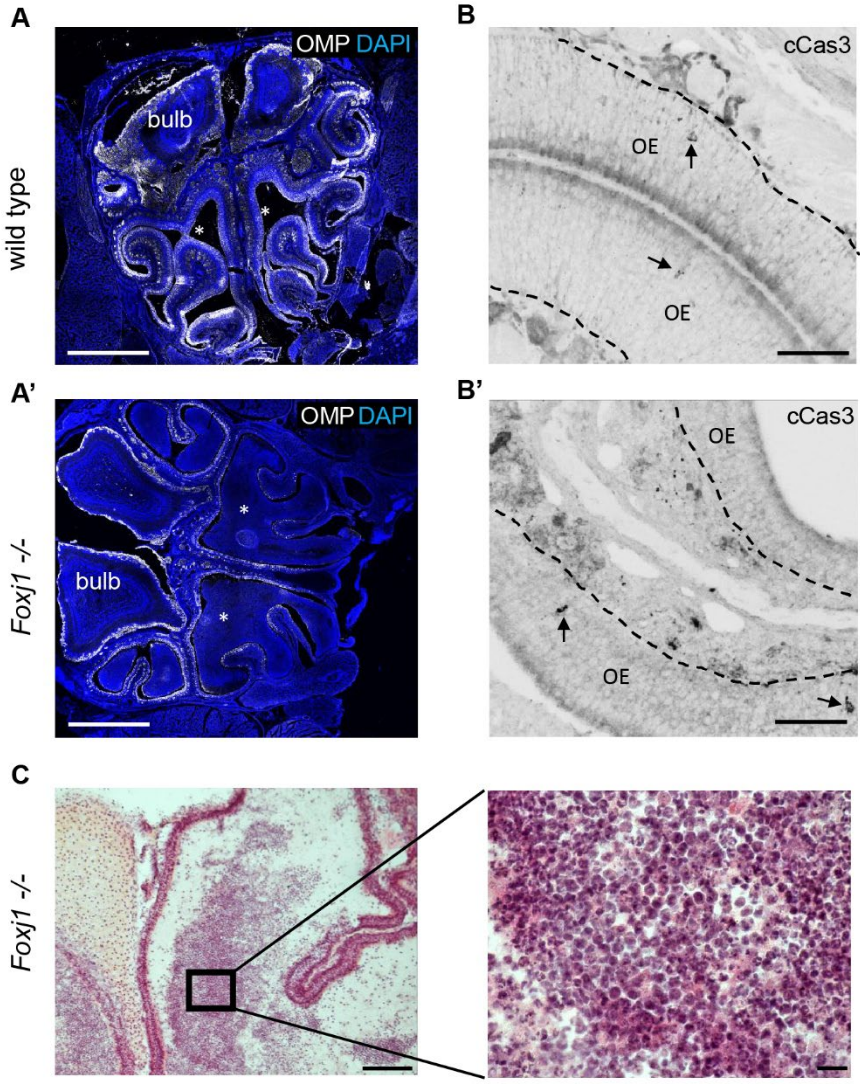


(A,B) Zoomed out view of the nasal cavity of the WT and *Foxj1-/-* mouse. Nasal cavity is denoted by asterisks.

(B,B') Level of apoptosis in the OE was similarly low in the WT (B) and *Foxj1-/-* (B') mouse (108±24, n=6, WT; 83±7 cells per $mm^{s}$, n=6, KO; p=0.554) as determined by immunostaining of cleaved Caspase 3 (arrows).

(C) Nasal cavity of the *Foxj1-/-* mouse contained neutrophiles as revealed by H&E stain.

Scale bar 1000 µm (A,A'), 100µm (B,B',C), and 10 µm (C inset).

**Figure S4: Transcriptional investigation of the olfactory epithelium across species.**

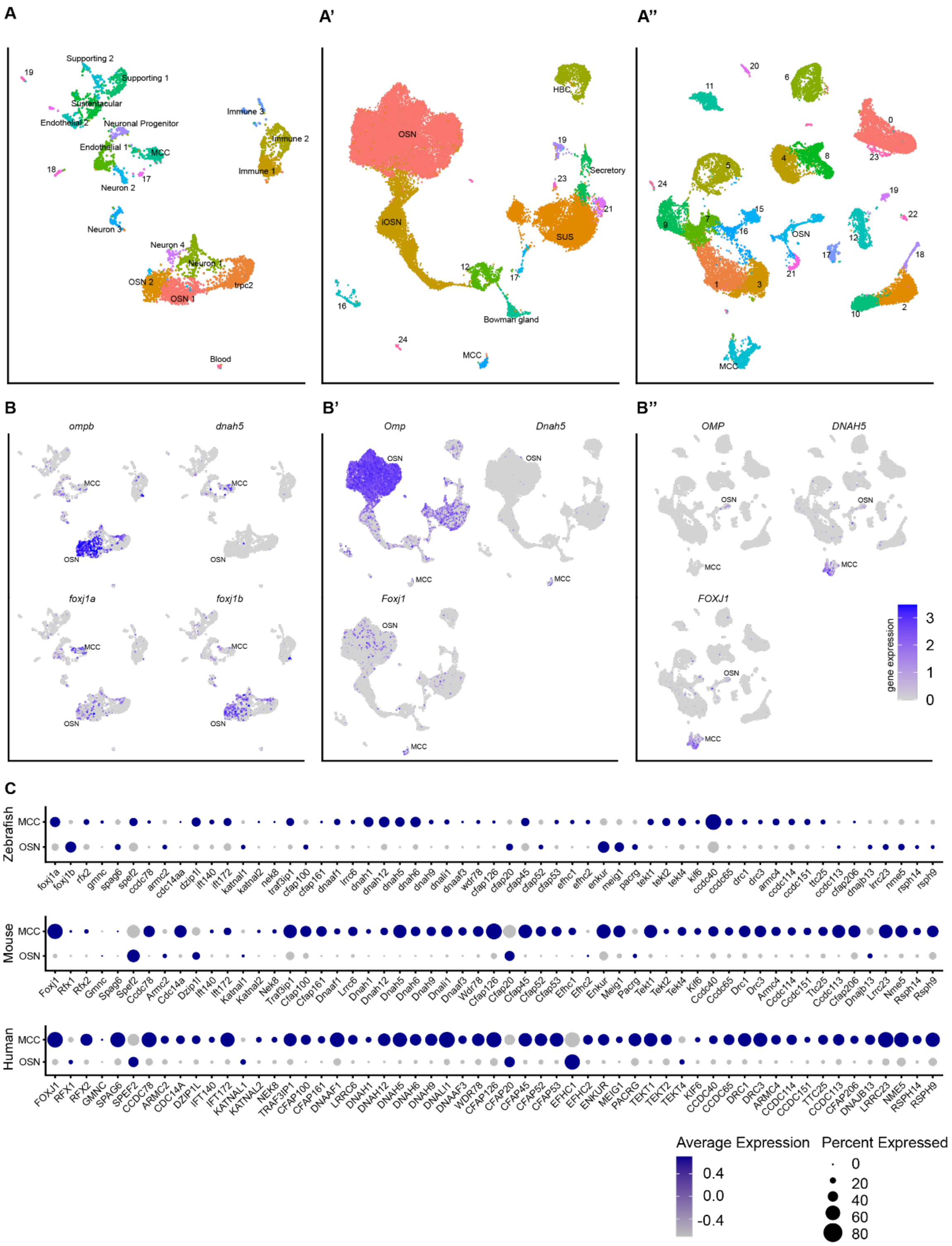


(A-A’’) Dimensionality reduction plots of detected cell clusters on single-cell RNA sequencing data in zebrafish, mouse and human olfactory epithelial tissue (calculated by umap). Cell clusters corresponding to MCCs and OSNs are shown.
(B-B’’) Marker genes *omp*, *dnah5* and *foxj1* expression across cell clusters.
(C) Dot-plot showing expression levels of *foxj1* target genes of interest in MCC and OSN cell clusters across species.

**Supplementary Video 1: The OSN cilia of the leopard frog *Rana pipiens* shows displays motility:**

An isolated OSN from the amphibian species *Rana pipiens* under the light-transmission microscopy. The movement of the OSN cilia can be observed. For details on how the cell suspension was achieved, see (Kleene and Gesteland 1991)
